# The mitophagy receptor BNIP3L/Nix coordinates nuclear calcium signaling to modulate the muscle phenotype

**DOI:** 10.1101/2023.03.18.532760

**Authors:** Jared T. Field, Donald Chapman, Yan Hai, Saeid Ghavami, Adrian R. West, Berkay Ozerklig, Ayesha Saleem, Julia Kline, Asher A. Mendelson, Jason Kindrachuk, Barbara Triggs-Raine, Joseph W. Gordon

**Affiliations:** Department of Human Anatomy and Cell Science, Rady Faculty of Health Sciences, University of Manitoba, Winnipeg, Canada; Department of Biochemistry and Medical Genetics, Rady Faculty of Health Sciences, University of Manitoba, Winnipeg, Canada; Department of Medical Microbiology & Infectious Diseases, Rady Faculty of Health Sciences, University of Manitoba, Winnipeg, Canada; Department of Physiology and Pathophysiology, Rady Faculty of Health Sciences, University of Manitoba, Winnipeg, Canada; Department of Internal Medicine, Rady Faculty of Health Sciences, University of Manitoba, Winnipeg, Canada; Department of Pediatrics and Child Health, Rady Faculty of Health Sciences, University of Manitoba, Winnipeg, Canada; Faculty of Kinesiology and Recreation Management, University of Manitoba, Winnipeg, Canada; Children’s Hospital Research Institute of Manitoba, Winnipeg, Canada

**Keywords:** BNIP3L (Nix), calcium signaling, mitophagy, muscle, myostatin

## Abstract

Mitochondrial quality control is critical in muscle to ensure both contractile and metabolic function. Nix is a BCL-2 family member, mitophagy receptor, and has been implicated in muscle atrophy. Human GWAS suggests altered Nix expression could predispose to manifestations of mitochondrial disease. To interrogate Nix function, we generated a muscle-specific knockout model. Nix knockout mice displayed a ragged-red fibre phenotype, along with accumulation of mitochondria and endo/sarcoplasmic reticulum with altered morphology. Intriguingly, Nix knockout mice were more insulin sensitive with a corresponding increase in glycogen-rich muscle fibres. Kinome- and gene expression analyses revealed that Nix knockout impairs NFAT and canonical myostatin signaling, with alterations in muscle fibre-type composition and evidence of regeneration. Mechanistic experiments demonstrated that Nix modulates mitophagy, along with ER-phagy leading to altered nuclear calcium signaling. Collectively, these observations identify novel roles for Nix coordinating selective autophagy, oxidative gene expression, and signaling pathways that maintain the muscle phenotype.

## Introduction

Mitochondrial quality control involves the fission of dysfunctional mitochondrial fragments, the removal of these fragments by PINK1/Parkin and receptor-mediated mitochondrial autophagy (ie. Mitophagy), combined with mitochondrial biogenesis and the fusion of nascent mitochondria with the existing mitochondrial network^1^. Aberrant mitochondrial quality control has been implicated in a number disease states, including muscle related pathologies, such as sarcopenia^2^, cachexia^3^, mitochondrial myopathies^4^, muscular dystrophies^5^, and the remodeling associated with obesity and insulin resistance^6^.

Nix (ie. BNIP3L) is a BCL-2 family member involved in the regulation of apoptosis, necrosis, macro-autophagy, and mitophagy^7^. In addition, Nix expression has recently been shown to be elevated during aging and has been implicated in starvation-induced muscle atrophy^8,9^. Human genome-wide association studies have identified non-coding single nucleotide polymorphisms within the Nix gene that increase the risk of early-onset schizophrenia and cognitive decline, both of which involve mitochondrial pathology, suggesting that reduced Nix expression may result in secondary mitochondrial myopathy^10,11^.

Nix is biologically active at both the mitochondria and the endoplasmic reticulum (ER)^7^. As a regulator of autophagy, Nix has been shown to activate macro-autophagy at the ER by disrupting the interaction between BCL-2 and Beclin-1, which has been proposed to take place at the IP3-receptor^7^. At the mitochondria, Nix acts as a selective autophagy receptor recruiting ATG8 proteins and the mTOR activator RHEB^12,13^, where deletion of both Nix and/or its homologue BNIP3 results in the age-related accumulation of large senescent mitochondria due to impaired mitochondrial pruning^14^. However, this model of Nix function may be incomplete, as BNIP3 can also promote selective ER-phagy (ie. Reticulophagy)^15^. Moreover, during a mitophagy response, we previously demonstrated that Nix promotes ER-dependent calcium signaling to activate the mitochondrial fission regulator DRP1, suggesting that both ER- and mitochondrial-localized Nix contribute to mitophagy^6^.

Given the diverse biological roles attributed to Nix, we interrogated Nix function in a novel muscle-specific knockout mouse with significant reduction in Nix expression in both red and white muscle, but not in the heart. This approach allowed us to assess the role of Nix in muscle biology without contaminant cardiovascular defects. Muscle-specific Nix knockout mice display a ragged-red fibre phenotype, along with accumulation of mitochondria and sarcoplasmic reticulum (SR) with altered morphology. Interestingly, Nix knockout also impaired NFAT and canonical myostatin (ie. GDF8) signaling resulting in alterations in muscle fibre-type composition. In addition, we present evidence of myopathy and regeneration in the absence of Nix. Mechanistic experiments in culture demonstrated that Nix is both necessary and sufficient to regulate mitophagy, but also ER-phagy through a distinct mechanism leading to nuclear calcium signaling. Collectively, these observations indicate that Nix has a biological role beyond autophagy/mitophagy, serving to modulate ER/SR homeostasis and signaling pathways that impact the muscle phenotype.

## Results

### Muscle-specific Nix knockout results in ragged red fibres, with the accumulation of mitochondria and sarcoplasmic reticulum with altered morphology

We used CRISPR/Cas9 technology to create a conditional allele with loxP sites flanking the second exon of the Nix (ie. *Bnip3l*) gene. Single-stranded donor DNAs were electroporated along with Cas9 and guide RNAs to insert loxP sites by homology directed repair (Fig.1A). Zygotes were implanted into pseudo-pregnant mice, and offspring were screened for insertion of loxP sites (Fig.1B). Mice containing both loxP sites were bred to homozygosity (Nix^fl/fl^) and crossed with human skeletal α-actin Cre mice (HSA-Cre). Cre-positive mice (Nix-HSA-KO) displayed reduced Nix expression in red, white, and mixed fibre-type muscles, but with no impact in the heart, while Cre expression alone did not influence Nix expression (Fig.1C-D). The degree of knockout in whole gastrocnemius was similar in magnitude to other reports using the HSA promoter^16^. Intriguingly, Gomori trichrome staining in Nix-HSA-KO mice revealed consistent evidence of ragged red muscle fibres in male mice by 10-weeks of age, which were notably absent in female mice, and mice of all other genotypes (Fig.1E; Supp Fig.1A). As the pathology of ragged red fibres has been associated with mitochondrial myopathy and the accumulation of subsarcolemmal mitochondria, we performed transition electron microscopy, which revealed large mitochondria both beneath the sarcolemma and amongst the myofibrils with altered morphology and cristae structure (Fig.1F; Supp Fig.1B, -C). Furthermore, we did not observe a compensatory increase in other mitophagy genes Parkin, BNIP3, or FUNDC1, but did observe a trend towards decreased expression (Supp Fig.1D). In addition, we observed the accumulation of SR membranes in close association with mitochondria (Supp Fig.1C). Intriguingly, removal of Nix from muscle had no significant impact on the macro-autophagy markers, LC3-II and P62 (Supp Fig.1E). To assess the impact of muscle-specific Nix knockout on metabolism, we performed metabolic caging, and observed a reduction in oxygen consumption, without changes in food consumption, body mass, or tibial length (Fig.1G; Supp Fig.1F). In addition, we evaluated oxygen consumption rate (OCR) in soleus muscle explants, which confirmed that Nix knockout reduced muscle oxygen consumption (Fig.1G). Nix-HSA-KO mice also displayed an increased resting respiratory exchange ratio (RER), suggesting increased reliance on carbohydrate metabolism, and a concurrent increase in blood lactate concentration (Fig.1H). Finally, we observed that Nix-HSA-KO mice had decreased running distance on a graded exercise treadmill test (Fig.1I). These observations implicate Nix in the regulation of both mitochondria and SR structure, and muscle function.

**Figure 1.**
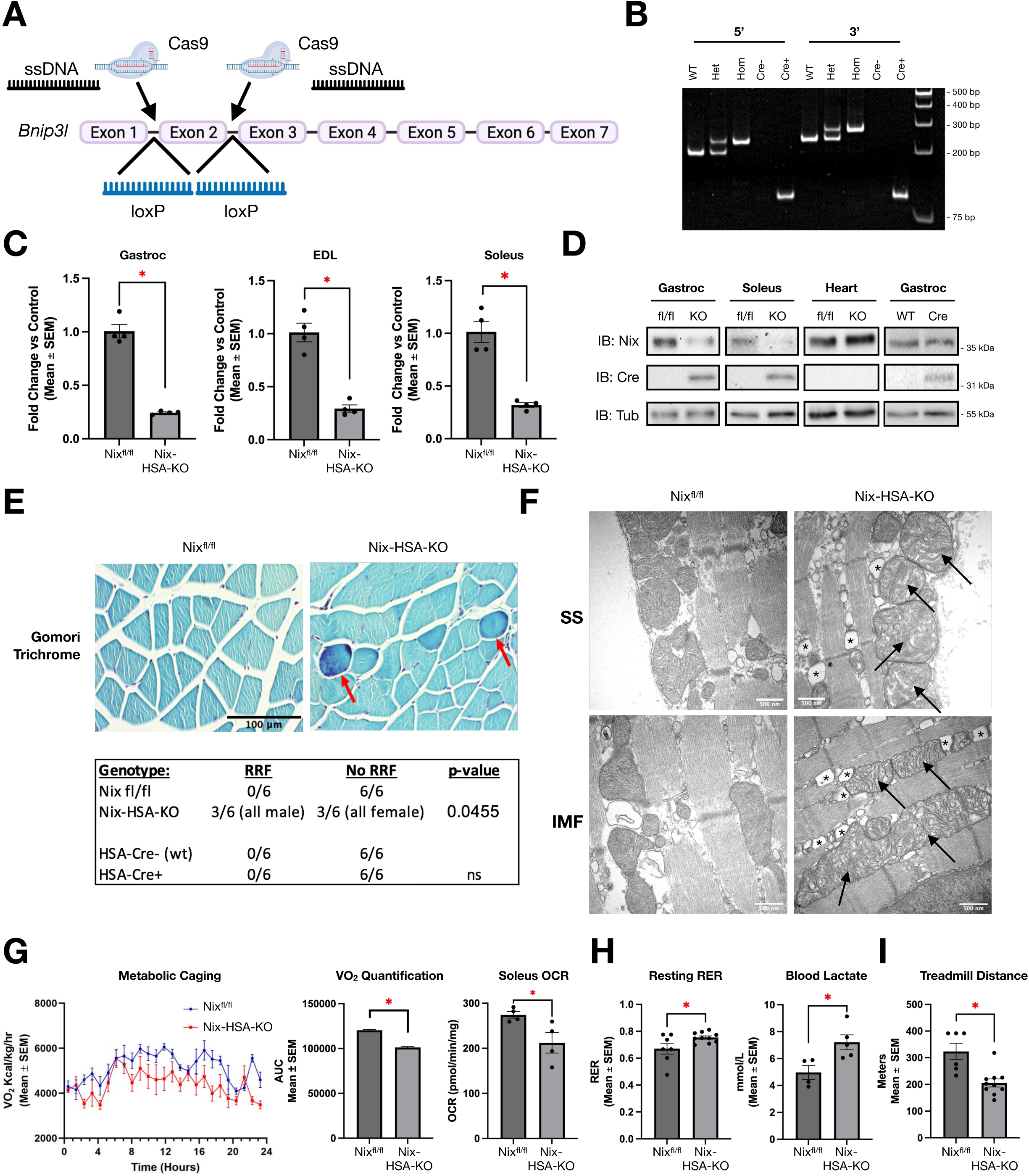
Signs of mitochondrial myopathy in muscle-specific Nix knockout mice. **A)** Schematic illustrating the generation of the floxed Nix (*Bnipl)* gene using CRISPR/Cas9. **B)** PCR detection of 5’ and 3’ loxP sites in heterozygous (Het) and homozygous (Hom; Nix^fl/fl^) mice and presence of the Cre transgene. **C)** Real-time PCR detection of Nix in Nix^fl/fl^ and knockout (Nix-HSA-KO) mice. Gastrocnemius/plantaris (Gastroc), extensor digitorum longus (EDL), and soleus muscle (n=4). **D)** Representative immunoblots of Nix and Cre by genotype. Wild-type (WT) and HSA-Cre (Cre). **E)** Gomori trichrome staining demonstrating ragged red fibres (red arrows) and quantification by genotypes (P-value determined by Chi-square test) in mice 10-12 weeks of age. **F)** Transmission electron microscopy of gastrocnemius muscle fibres in longitudinal section showing morphology of subsarcolemmal (SS) and intermyofibrillar (IMF) mitochondria. Abnormal mitochondrial morphology (black arrows) and sarcoplasmic reticulum membranes (*). **G)** Oxygen consumption and area under the curve quantification of Nix^fl/fl^ and Nix-HSA-KO male mice 10-12 weeks of age (n=4; left). Oxygen consumption rate (OCR) of soleus muscle ex plants using a Seahorse XFe24 analyzer (right). **H)** Respiratory exchange ratio (RER) calculated from resting metabolic cage data (left) and resting non-fasting blood lactate (right). **I)** Maximal distance ran in graded exercise treadmill test (n=6-10). Data shown as mean with error bars indicating standard error of the mean. *p<0.05.

### Kinome analysis of muscle-specific Nix knockout mice reveals increased signaling responses involved in nutrient storage and impaired calcium and TGF-β signaling

Previously, we demonstrated that Nix orchestrates both ER-to-mitochondrial calcium transfer and mTOR signaling during a mitophagy response^6,17^. As ER calcium depletion, ER-stress, and ER-phagy often occur concurrently, we hypothesized Nix-HSA-KO mice may present with defects in calcium signaling. To evaluate this hypothesis *in vivo*, we performed kinome microarray analysis using the workflow illustrated in Figure 2A, and described previously^18,19^. Pathway overrepresentation and Gene Ontology analyses were performed using InnateDB (Fig.2A; Supp. Fig.2A, B), which identified upregulation of pathways involved in lipid and glycogen biosynthesis, and downregulation of pathways involved with TGF-β and calcium signaling. Inspection of individual signaling pathways, revealed altered phosphorylation of key proteins involved in the insulin signaling pathway and decreased phosphorylation of phosphorylase b kinase (KPB1) involved in glycogen breakdown (Fig.2B). To evaluate the physiological significance of altered insulin signaling, we performed insulin tolerance tests in fasted Nix^fl/fl^ and Nix-HSA-KO mice, which confirmed enhanced insulin sensitivity in the Nix-HSA-KO mice (Fig.2C; Supp Fig.2C). In addition, Periodic acid– Schiff (PAS) staining of gastrocnemius/plantaris muscle identified an increased number of glycogen-rich muscle fibres in Nix-HSA-KO mice (Fig.2D, Supp Fig.2C). We also performed kinome microarray analysis in muscle tissue following administration of insulin. InnateDB and Gene Ontology revealed overrepresentation of insulin activated pathways, including Rho-RAC1 which are involved in GLUT4 translocation, and several other pathways involved in cell growth and metabolism in the Nix-HSA-KO mice (Supp Fig.2D).

**Figure 2.**
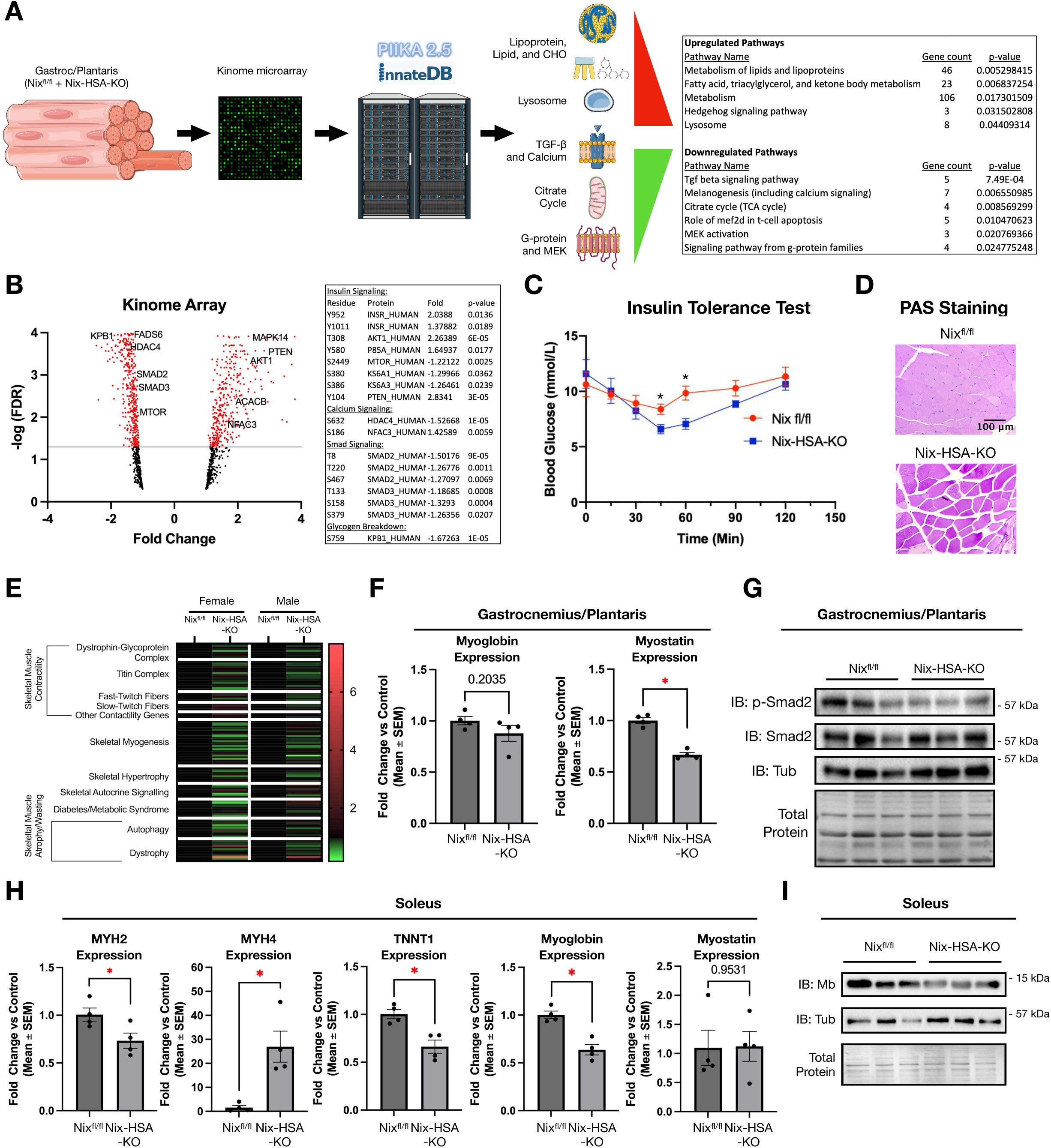
Deletion of Nix alters multiple cell signalling cascades in muscle. **A)** Kinomics analysis workflow and pathway analysis by InnateDB. **B)** Volcano plot of individual phospho-residues showing fold change in Nix-HSA-KO mice over Nix^fl/fl^ male mice at 10-12 weeks of age. P-values were adjusted using the false discover rate method (FDR)(n=3). Table of phospho-residues involved in insulin, calcium, and TGF-β/SMAD signaling as well as glycogen breakdown. **C)** Insulin tolerance test in fasted Nix^fl/fl^ and Nix-HSA-KO male mice (n=5). **D)** Periodic acid Schiff (PAS) staining for glycogen content in gastrocnemius/plantaris muscle of male mice. **E)** Heat map generated from myogenesis and myopathy qPCR array. Green, red, and black colours indicate down-, up-regulation, or no change respectively (n=4). **F)** Gene expression of myoglobin and myostatin in gastrocnemius/plantaris muscle of male mice (n=4). **G)** Immunoblot of total and phospho-Smad2 expression in extracts from gastrocnemius/plantaris muscle (Tub=β-tubulin) (n=3). **H)** Gene expression of MYH2, MYH4, TNNT1, myoglobin and myostatin in soleus muscle extracts from male mice (n=4). **I)** Immunoblot of myoglobin (Mb) expression in soleus muscle extracts (n=3). Data shown as mean with error bars indicating standard error of the mean. *p<0.05.

Our kinome analysis also identified reduced phosphorylation of HDAC4 at Ser-632, and increased phosphorylation of NFATc3 (NFAC3) at Ser-186 (Fig.2B). As these residues have been previously implicated in calcium-dependent gene expression^20^, we performed multiple PCR-based arrays targeting myogenesis and myopathy genes comparing male and female mice and red and white muscle groups in Nix^fl/fl^ and Nix-HSA-KO mice (Fig.2E; Supp Table 1). Interestingly, female knockout mice displayed a greater number of genes with reduced expression compared to male knockout mice, despite the absence of ragged red fibres. However, we observed that female mice also had reduced expression of PGC-1α and increased myogenin expression, which may protect against this phenotype (Supp Fig.2E). We also examined the expression of genes involved in the unfolded protein response (UPR) and did not observe significant changes, although there was a trend towards reduced GRP94 expression in Nix-HSA-KO mice (Supp Fig.2F). Examination of two known NFAT-target genes, myoglobin and myostatin^20,21^, in gastrocnemius/plantaris and soleus muscle of male mice revealed that myostatin expression was reduced in gastrocnemius/plantaris, while myoglobin expression was reduced in soleus muscle (Fig.2F, -H). This suggests some degree of muscle-group or fibre-type specific effect of Nix knockout, which is supported by the data in Supplemental Table 1 comparing gene expression in different muscle groups. As myostatin is a member of the TGF-β family that signals through Smad2/3, we confirmed that Nix knockout reduced p-Smad2 in gastrocnemius/plantaris (Fig.2G Supp Fig.3A), consistent with the kinome data (Fig.2B). Next, we examined the expression of other muscle oxidative genes in soleus muscle and observed reductions in MYH2 (Type IIa; fast oxidative) and TNNT1 (Troponin-T; slow muscle) (Fig.2H). Moreover, we observed a marked increase in MHY4 (Type IIb; fast glycolytic) expression. Finally, we confirmed the decreased expression of myoglobin in the soleus of Nix-HSA-KO mice by western blot (Fig.2I; Supp Fig.3B). Collectively, these observations suggest that Nix knockout in muscle enhances insulin signaling to promote nutrient storage and impairs calcium-dependent gene expression and downstream myostatin activity.

**Figure 3.**
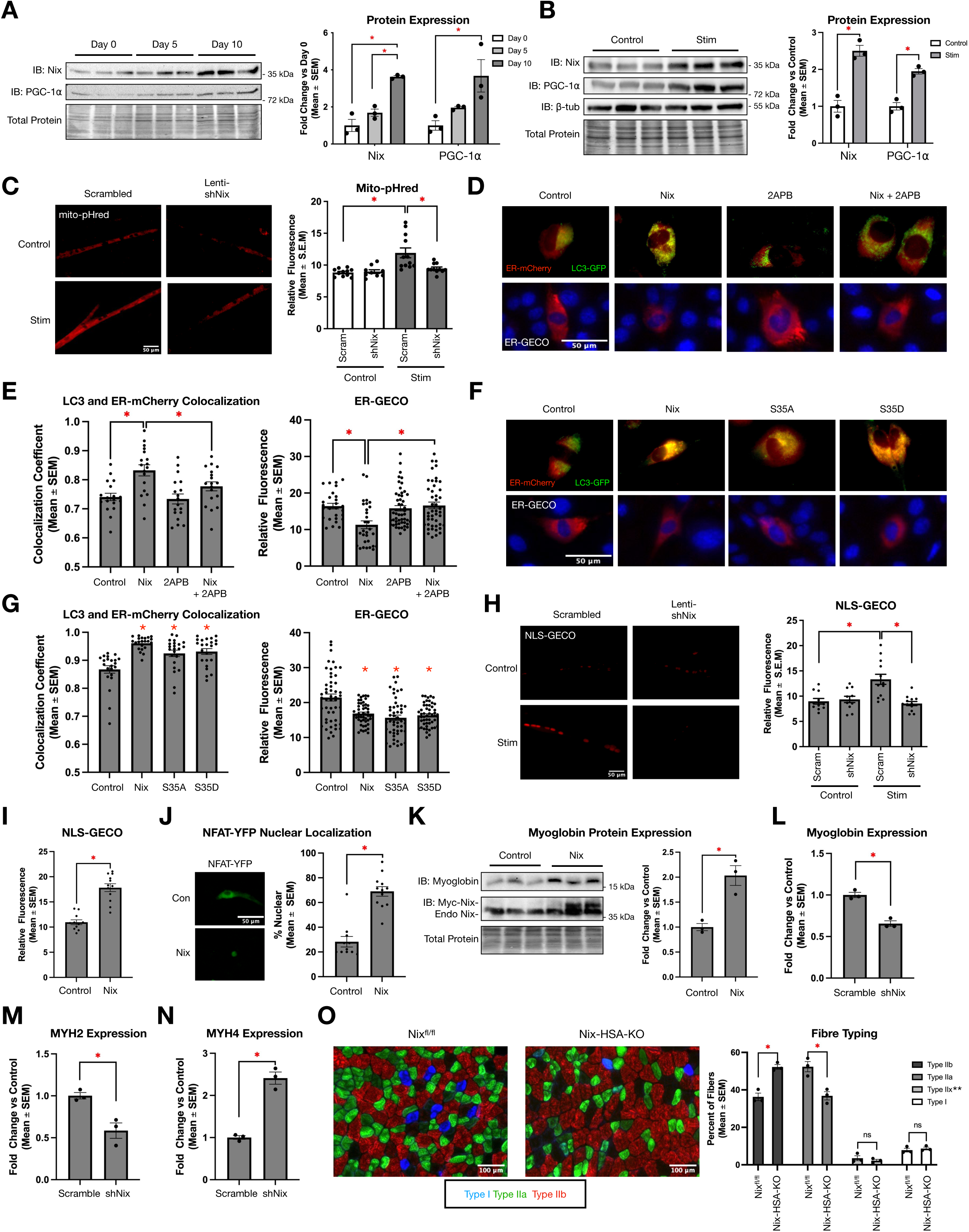
Nix is both necessary and sufficient to activate nuclear calcium signaling and gene expression. A-B) Immunoblots of Nix and PGC-1α and quantification of protein levels in differentiating C2C12 myotubes (a) and in electrically paced (Stim) C2C12 myotubes (b)(n=3). **C)** Representative images and quantification of the mitophagy biosensor mito-pHred in myotubes with or without electrical pacing and knockdown of Nix with lentriviral RNA interference (Lenti-shNix). **D-E)** C2C12 cells transfected with ER-mCherry, LC3-GFP, and Nix, and treated with 2APB (2µM), as indicated (above) or ER-GECO (below). Quantification performed by Pearson’s colocalization coefficient or mean fluorescence. **F-G)** C2C12 cells were transfected as in (d) with Nix, Nix-S35A, or Nix-S35D, and quantified as in (e). **H)** Representative images and quantification of the nuclear calcium biosensor (NLS-GECO) in paced myotubes with and without Lenti-shNix. **I)** NLS-GECO in C2C12 cells transfected with empty vector or Nix. **J)** Representative images and quantification of sub-cellular distribution of NFAT-YFP in C2C12 cells expressing Nix or empty vector. **K)** Immunoblot and quantification of myoglobin expression in C2C12 cells transfected with Myc-Nix (n=3). Endo-Nix indicates endogenous Nix. **L-N)** C2C12 cells were transfected with an shRNA targeting Nix (shNix) or scrambled control (Scramble) and allowed to differentiate for 6-days. RNA expression of myoglobin (l) MYH2 (m) and MYH4 (n) in C2C12 myotubes. **O)** Immunofluorescence detection of specific muscle fibre-types in the central region of gastrocnemius/plantaris muscle of male mice 10-12 weeks of age. Type IIa (green; MYH2), type IIb (red; MYH4), and type I (blue; MYH7). **The percentage of type IIx fibres was estimated by non-staining regions. Data shown as mean with error bars indicating standard error of the mean. *p<0.05.

### Nix is both necessary and sufficient to activate nuclear calcium signaling and gene expression

Using two independent C2C12 models of mitochondrial biogenesis, differentiation and electrical pacing, we observed that Nix expression increased in parallel with PGC-1α (Fig.3A, -B). These observations suggest that quality control pathways are activated concurrently with mitochondrial biogenesis. To evaluate the role of Nix in this hypothesis, we utilized the mitophagy biosensor mito-pHred^6^, in combination with a validated lentiviral shRNA targeting Nix (Lenti-shNix; Supp Fig.3C)^6^ and electrical pacing. Shown in Figure 3C, electrical pacing (Stim) increased mito-pHred fluorescence, which was prevented by Nix knockdown. As Nix-HSA-KO mice display evidence of ER/SR membrane accumulation, we speculated that Nix may also be involved in selective removal of endoplasmic reticulum by ER-phagy. Thus, we transfected C2C12 cells with ER-targeted mCherry (ER-mCherry) and LC3-GFP to monitor ATG8 recruitment to the ER in the presence of Nix, which represents an early event in ER-phagy. We observed that Nix significantly increases the colocalization of LC3-GFP with ER-mCherry, but this colocalization was blocked by the IP3-receptor antagonist 2APB (Fig.3D-E). In addition, we monitored ER calcium depletion in a parallel series of experiments using the ER-targeted calcium biosensor ER-GECO, which identified that ER calcium levels and ER-phagy have an inverse relationship (Fig.3D-E). Nix also induced a similar degree of colocalization between ER-Emerald and Lysotracker, a late indicator of the autophagy process (Supp. Fig.D). As an additional control, cells were treated with the lysosomal inhibitor bafilomycin A1, which also increased LC3 colocalization with ER-mCherry (Supp. Fig.3E). As phosphorylation of Nix at Ser-35 has been previously shown to enhance LC3/ATG8 recruitment to mitochondria during mitophagy^12,22^, we evaluated the effect of neutral and phospho-mimetic mutations of Nix at this residue (ie. S35A and S35D, respectively). Intriguingly, wild-type Nix, S35A, and S35D all had a similar impact on both LC3-GFP recruitment to the ER and ER calcium depletion (Fig.3F-G); however, when we evaluated colocalization between LC3-GFP and mito-mCherry, Nix-S35D induced a greater degree of LC3 recruitment to the mitochondrial than Nix-S35A (Supp. Fig.3F). Collectively, these data demonstrate that ER calcium depletion is a key mechanism by which Nix induces ER-phagy, and that phosphorylation at Ser-35 is more important in recruitment of LC3 to mitochondria.

Next, we evaluated the effect of Nix on down-stream calcium signaling. We transfected C2C12 cells with the nuclear-targeted calcium biosensor, NLS-GECO^23^, followed by differentiation and electrical pacing. Intriguingly, pacing increased nuclear calcium, but was attenuated by knockdown with Lenti-shNix (Fig.3H). In addition, transfection of Nix into C2C12 cells increased NLS-GECO fluorescence, which was inhibited by the IP3-receptor antagonist 2ABP (Fig.3I; Supp Fig.3G). Moreover, cell fractionation studies revealed that Nix preferentially accumulates at the ER/SR following pacing (Supp Fig.3H). These observations identify a dual role for Nix, regulating selective autophagy and nuclear calcium signalling. To evaluate if increased nuclear calcium induced by Nix translates into changes in gene expression, we first transfected NFAT-YFP with and without Nix in C2C12 cells, and observed that Nix expression was sufficient to increase the nuclear accumulation of NFAT-YFP (Fig.3J). Nix expression also increased the endogenous expression of the NFAT-target gene myoglobin (Fig.3K). Next, we returned to the C2C12 differentiation model, but knocked-down Nix using a plasmid-based shRNA prior to differentiation (Supp Fig.3I)^6,24^, and evaluated myoglobin expression, and the myosin heavy chain genes that were altered *in vivo*. Consistent, with the data in Figure 2H, shNix decreased myoglobin and MYH2 expression, but increased MYH4 expression (Fig.3L-N). As calcium and NFAT signaling have been previously shown to alter the muscle fibre-type composition, promoting oxidative fibre-types^20^, we evaluated fibre-type composition in Nix^fl/fl^ and Nix-HSA-KO mice. Shown in Figure 3O, Nix-HSA-KO mice have a reduced number of oxidative type IIa fibres and an increase in glycolytic type IIb fibres in the gastrocnemius/plantaris muscle groups, without changes in type I or IIx fibres. Similar results were observed in soleus muscle, where Nix-HSA-KO mice have decreased numbers of oxidative fibres (Type I) and increased numbers of glycolytic fibres (Type IIb and IIx; Supp. Fig.3J).

### Muscle-specific Nix knockout results in a myopathy with central nuclei and regeneration, without an increase in muscle fibrosis

Our Gene Ontology enrichment analysis in Nix-HSA-KO mice identified biological responses involved with proteolysis, cell growth, and phospholipid and steroid biosynthesis, all of which have been implicated in muscle repair and regeneration (Supp Fig.2B). Consistent with this, muscle cross-sections from Nix-HSA-KO display central nuclei (Fig.4A), and we observed increased expression of embryonic myosin heavy chain (MYH3; Fig.2B). In addition, examination of gene expression identified increased Pax7 expression, with a corresponding increase in the satellite cell mitogen TNF (Fig.4C)^25^, which can operate in a reciprocal manner to myostatin during muscle regeneration. Interestingly, we also observed decreased expression of dystrophin (DMD) and increased expression of caspase 3 (Casp3), known for its role in apoptosis but also as a regulator of Pax7 function (Fig.4C)^26^. The expression of NRF2, a transcription factor involved in the regulation of antioxidant gene expression and modulation of satellite cell function, was also increased in Nix-HSA-KO mice (Fig.4D)^27^. We also examined signaling pathways identified through our kinome analysis and observed tyrosine phosphorylation of several receptors implicated in muscle regeneration, including INAR1, EPHB2, PDGFRA, and FGFR1 (Fig.4E). Our kinome analysis also identified phosphorylation of p38α (MK14) at the inhibitory Thr-123 residue, consistent with an established role of p38α in the repression of Pax7 expression^28,29^. Finally, we observed an increased number of Pax7 positive satellite cells within gastrocnemius/plantaris muscle in Nix-HSA-KO mice, without alterations in muscle fibrosis (Fig.4F, -G). Collectively, these observations demonstrate that muscle-specific Nix knockout results in a myopathy with compensatory regeneration.

**Figure 4.**
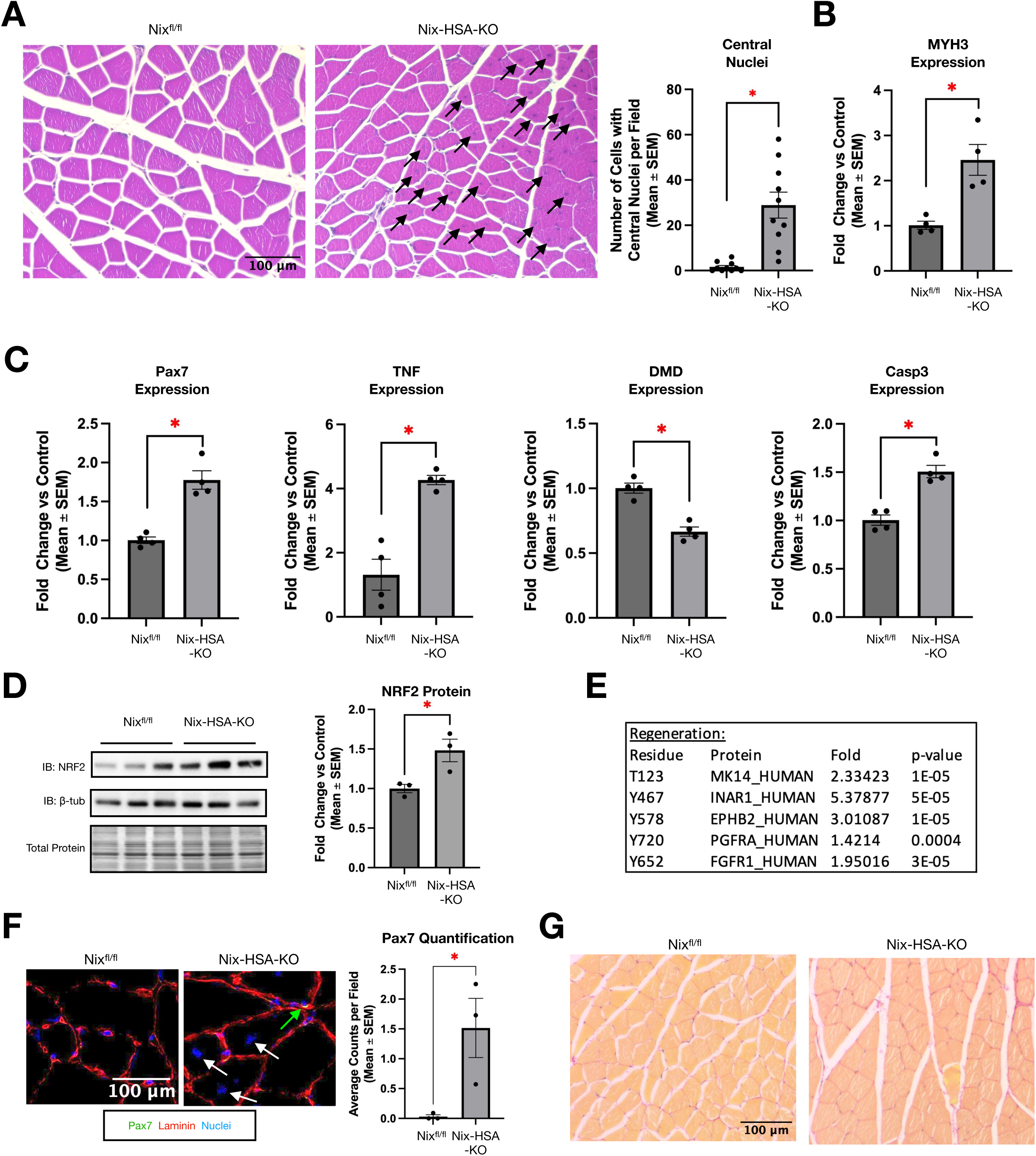
Muscle-specific Nix knockout results in a myopathy with central nuclei and regeneration. **A)** H&E staining of gastrocnemius/plantaris muscle from Nix^fl/fl^ and Nix-HSA-KO male mice showing presence of centrally located nuclei (black arrows), and quantification in male mice 10-12 weeks of age. **B)** Gene expression of embryonic myosin heavy chain (MYH3), an indicator of regeneration (n=4). **C)** Gene expression of markers of muscle satellite cell activation in male mice. TNF=TNFα, DMD=dystrophin, Casp3=caspase-3 (n=4). **D)** Immunoblot and quantification of NRF2 protein (n=3). **E)** Kinome analysis of phospho-residues involved in muscle regeneration and/or satellite cell activation (n=3). **F)** Immunofluorescence and quantification of Pax7 positive cells (green with green arrow), laminin (red) in gastrocnemius/plantaris muscle (nuclei counterstained blue). Central nuclei identified by white arrows (n=3). **G)** Picrosirius red staining of gastrocnemius/plantaris muscle demonstrating no overt fibrosis in Nix-HSA-KO male mice 10-12 weeks of age (connective tissue stains red). Data shown as mean with error bars indicating standard error of the mean. *p<0.05.

## Discussion

Emerging GWAS evidence suggests that polymorphisms within the Nix gene are associated with mitochondrial related pathology^10,11^. Although the precise impact of these polymorphisms is not currently known, Nix has been experimentally implicated in muscle atrophy, aging, lipotoxicity, and cardiac remodeling^6,8,9,24,30^, where mitochondrial dysfunction has been shown to contribute to the pathogenesis of these disorders. In this report, we interrogate Nix function in muscle using a novel mouse knockout model, which present with evidence of compensated mitochondrial myopathy. Moreover, this mouse model has uncovered novel aspects of Nix function, that could be of translational importance as skeletal muscle is often used as diagnostic proxy in the evaluation of neuropathology. The skeletal muscle phenotype has been described in terms of distinct muscle fibre-types that are commonly identified by the expression of a specific myosin heavy chain gene^20^. Our data identifies Nix, a known regulator of mitophagy^6,14^, as a modulator of muscle fibre-type composition by regulating several signaling pathways that target gene expression. Our observations suggest Nix-induced mitophagy parallels mitochondrial biogenesis to mechanistically couple mitochondrial turn-over with oxidative gene expression, ultimately modifying the muscle phenotype.

Muscle-specific Nix knockout also results in the accumulation of ER/SR membranes, while mechanistic experiments demonstrate that Nix modulates ER-phagy and calcium homeostasis. Interestingly, we observed that ER calcium depletion is an important mechanism triggering ER-phagy, which is distinct from Nix-induced mitophagy which depends on direct ATG8/LC3 recruitment^12,22,31^. We also observed sex-specific differences in the present study, notably the absence of ragged red fibres in female mice. Interestingly, female mice displayed differential regulation of gene expression, including reduced PGC-1α expression. Potentially, this prevents mitochondrial accumulation and averts the ragged red fibre phenotype. In addition, we observed increased myogenin expression in female mice, and increased IL-1β in female knockout mice, which may contribute to increased muscle regeneration that prevented the appearance of ragged red fibres.

The present study also confirms and extends our previous work that identified Nix as a modulator of mTORC1 in the regulation of insulin signaling^6,24^. Previously, we demonstrated that Nix is responsive to diacylglyceride accumulation, known to promote insulin resistance^32^, to activate mitophagy and inhibit insulin signaling through a mechanism contingent on IRS-1 phosphorylation^6,24^. In the present study, we observed alterations in signaling pathways associated with glycogen and lipid metabolism in Nix-HSA-KO mice, which may be a direct effect of Nix deletion, or secondary to changes in the fibre-type. Future work should elucidate the role of Nix in muscle during conditions of altered mitochondrial biogenesis, such as exercise training, denervation, and sarcopenia.

Perhaps the most intriguing observation in muscle-specific Nix knockout mice is the alterations in TGF-β/myostatin signaling, without altering expression of other secreted metabolic regulators, such as FGF21 or GDF15 (Supp Table 1). Elevated myostatin secretion has been implicated in muscle atrophy, insulin resistance, satellite cell inhibition, activation of fibro-adipogenic progenitors (FAPs), and the repression of muscle genes, such as MYH4^33–35^. Thus, the reduction in myostatin expression and impaired Smad2 signaling in Nix-HSA-KO mice is consistent with the phenotypic alterations, including increased MYH4 expression, increased insulin sensitivity, and regeneration without overt fibrosis.

In summary, our results identify a new biological role for Nix maintaining mitochondrial, endoplasmic reticulum and calcium homeostasis, ultimately modulating the oxidative muscle phenotype. Selective targeting of specific Nix functions could alleviate many of the detrimental manifestations of muscle and metabolic disease.

## Materials and Methods

### Generation of Muscle-Specific Deletion of Nix in Mice

All procedures were approved by the Animal Care Committee of the University of Manitoba, which adheres to the principles developed by the Canadian Council on Animal Care (CCAC). Nix^fl/fl^ mice were designed and generated by the University of Manitoba Transgenic Facility using the IDT Alt-R^TM^ CRISPR-Cas9 system together with a ssDNA donors in C57BL/6N zygotes to sequentially insert loxP sites flanking exon 2 of the *Bnip3l* (*Nix*) gene. Guide RNAs, 5’-ggaactatttgagcgctttg-3’ (5’ side) and 5’-ttggttgacccgtttcatcc-3’ (3’ side) together with donors containing the loxP site and 60 bp arms matching the sequence upstream and downstream of the desired insertion site were purchased from Integrated DNA Technologies, Inc (USA). Removal of exon 2 creates a premature stop codon in the Nix transcript that would be expected to result in nonsense mediated decay. This strain is available at The Jackson Laboratory Repository with the JAX Stock No. 039220. Hemizygous HSA-Cre mice were obtained from Jackson Labs (ACTA1-cre, #006149) and crossed with Nix^fl/fl^ mice to conditionally ablate Nix specifically in skeletal muscle (Nix-HSA-KO). Experimentation was carried out on 10-12 week-old mice.

### Physiological Assays

Insulin response was characterized by insulin tolerance test using intraperitoneal injection of bovine insulin (0.56 IU/g of body weight; Sigma I0516). Exercise tolerance was assessed by treadmill running mice to exhaustion while measuring distance. Baseline metabolism measurements were performed by indirect calorimetry using metabolic cages (Columbus Instruments CLAMS) over a period of 24 hours after a familiarization period (24 hours). Resting respiratory exchange ratio (RER) was calculated from individual data collected in a metabolic cage during the light cycle. Blood lactate was measured at rest from a left ventricle cardiac puncture using an EPOC Blood Analysis System (Siemens Healthcare Limited). Oxygen consumption rate (OCR) was evaluated using a Seahorse XFe24 Analyzer (Agilent) using intact soleus muscle ex plants^36^.

### Histology, Immunofluorescence, and Electron Microscopy

Conventional histological stains (H&E, PAS, Picrosirius Red, Gomori Trichrome) were performed on formalin-fixed sections of muscle following standard protocols in the University of Manitoba Histology and Electron Microscopy core facility. Immunofluorescence experiments were performed on fresh-frozen sections of muscle using antibodies listed in the supplemental methods. Transmission electron microscopy was performed in longitudinal sections of muscle fixed in glutaraldehyde and analysed by an expert pathologist blinded to the experimental conditions^37^.

### Cell Culture, Electrical Pulse Stimulation, and Transfection

C2C12 myoblasts were cultured as described previously^6^. C2C12 cells were differentiated into myotubes for up to 10 days in DMEM (without pyruvate) with 0.5% FBS and supplemented with ITS (Gibco 41400-045), as indicated. Ten-day differentiated myotubes were paced by electrical pulse stimulus (12V, 1Hz, 1hr; Ion Optix C-Pace EM). Cells were transfected with plasmids (see supplemental methods) using jetPRIME transfection reagent (Polyplus). The Nix plasmids are available through Addgene (plasmid # 100795, #100770, #197562, #197563)^17^. Serine-35 mutations were constructed using the Q5 Site-Directed Mutagenesis Kit (New England Biolabs). Primers are listed in the supplemental methods.

### Real Time qPCR, Immunoblotting, and Kinome Analysis

RNA and protein were isolated from muscle and cells^6^. Array-based qPCR (BioRad, myogenesis and myopathy, SAB gene list) used the built-in primers, while conventional qPCR was performed using primers listed in the supplemental methods. Immunoblot analysis of proteins was performed by SDS-PAGE followed by immunoblotting with antibodies listed in the supplemental methods. Kinome analysis was performed on protein lysates, as previously described^18,19^.

### Statistical Analyses

Statistical analyses were performed using GraphPad Prism software. Unpaired 2-tailed t-test, Chi square, one-way ANOVA (post-hoc: Bonferroni), and two-way ANOVA (post-hoc: Tukey) were used to evaluate significance (α =0.05).

## Abbreviations

ER: Endoplasmic reticulum;
SR: Sarcoplasmic reticulum

## Acknowledgements

This work was support by the Natural Science and Engineering Research Council (NSERC) Canada through a Discovery Grant, and a Diabetes Canada-End Diabetes Grant to JWG. Seed funding was provided by the Children’s Hospital Research Institute of Manitoba, the DREAM research theme, and the Manitoba Centre for Nursing and Health Research. Transgenic and histological services were subsidized by the University of Manitoba Rady Faculty of Health Sciences. J.K. is supported by a Tier 2 Canada Research Chair provided by the Canadian Institutes of Health Research (950-231498; CRC-2021-00098). J.T.F. is supported by an Alexander Graham Bell studentship from NSERC Canada. The authors wish to acknowledge Xiaoli Wu for technical assistance generating Nix^fl/fl^ mice, Farhana Begum for histological support, Andrew Tse for confocal assistance, Dr. David Sontag for blood analysis support, Dr. Richard LeDuc for helpful bioinformatics discussion, and Dr. James Thliveris for electron microscopy support. Images from Bioicons and BioRender were used in the generation figures.

## Author Contributions

J.T.F., J.K., B.T-R., and J.W.G designed research; J.T.F., D.C., S.G., A.R.W., A.S., J.K., A.M., Y.H., and J.K. performed research; J.T.F., J.K., A.M., B.T-R., and J.W.G. analyzed data; and J.T.F., J.K., B.T-R., and J.W.G. wrote the paper.

## Competing Interest Statement

The authors have no competing interests to disclose.

## Notes

### Competing Interest Statement

The authors have declared no competing interest.

### Summary of Updates

Additional data including blood lactate and RER, along with some supplemental information.

## References

1. Shirihai OS, Song M, Dorn GW. How Mitochondrial Dynamism Orchestrates Mitophagy. Circulation Research 2015; 116:1835–49.

2. Triolo M, Hood DA. Manifestations of Age on Autophagy, Mitophagy and Lysosomes in Skeletal Muscle. Cells 2021; 10:1054.

3. VanderVeen BN, Fix DK, Carson JA. Disrupted Skeletal Muscle Mitochondrial Dynamics, Mitophagy, and Biogenesis during Cancer Cachexia: A Role for Inflammation. Oxid Med Cell Longev 2017; 2017:3292087.

4. Mito T, Vincent AE, Faitg J, Taylor RW, Khan NA, McWilliams TG, Suomalainen A. Mosaic dysfunction of mitophagy in mitochondrial muscle disease. Cell Metab 2022; 34:197–208.e5.

5. Reid AL, Alexander MS. The Interplay of Mitophagy and Inflammation in Duchenne Muscular Dystrophy. Life (Basel) 2021; 11:648.

6. da Silva Rosa SC, Martens MD, Field JT, Nguyen L, Kereliuk SM, Hai Y, Chapman D, Diehl-Jones W, Aliani M, West AR, et al. BNIP3L/Nix-induced mitochondrial fission, mitophagy, and impaired myocyte glucose uptake are abrogated by PRKA/PKA phosphorylation. Autophagy 2021; 17:2257–72.

7. Field JT, Gordon JW. BNIP3 and Nix: Atypical regulators of cell fate. Biochimica et Biophysica Acta (BBA) - Molecular Cell Research 2022; 1869:119325.

8. Mammucari C, Milan G, Romanello V, Masiero E, Rudolf R, Del Piccolo P, Burden SJ, Di Lisi R, Sandri C, Zhao J, et al. FoxO3 controls autophagy in skeletal muscle in vivo. Cell Metab 2007; 6:458–71.

9. Abadir P, Ko F, Marx R, Powell L, Kieserman E, Yang H, Walston J. Co-Localization of Macrophage Inhibitory Factor and Nix in Skeletal Muscle of the Aged Male Interleukin 10 Null Mouse. J Frailty Aging 2017; 6:118–21.

10. Li QS, Parrado AR, Samtani MN, Narayan VA, Alzheimer’s Disease Neuroimaging Initiative. Variations in the FRA10AC1 Fragile Site and 15q21 Are Associated with Cerebrospinal Fluid Aβ1-42 Level. PLoS One 2015; 10:e0134000.

11. Lam M, Chen C-Y, Li Z, Martin AR, Bryois J, Ma X, Gaspar H, Ikeda M, Benyamin B, Brown BC, et al. Comparative genetic architectures of schizophrenia in East Asian and European populations. Nat Genet 2019; 51:1670–8.

12. Rogov VV, Suzuki H, Marinković M, Lang V, Kato R, Kawasaki M, Buljubašić M, Šprung M, Rogova N, Wakatsuki S, et al. Phosphorylation of the mitochondrial autophagy receptor Nix enhances its interaction with LC3 proteins. Sci Rep 2017; 7:1131.

13. Melser S, Chatelain EH, Lavie J, Mahfouf W, Jose C, Obre E, Goorden S, Priault M, Elgersma Y, Rezvani HR, et al. Rheb regulates mitophagy induced by mitochondrial energetic status. Cell Metab 2013; 17:719–30.

14. Dorn GW. Mitochondrial Pruning by Nix and BNip3: An Essential Function for Cardiac-Expressed Death Factors. J of Cardiovasc Trans Res 2010; 3:374–83.

15. Hanna RA, Quinsay MN, Orogo AM, Giang K, Rikka S, Gustafsson ÅB. Microtubule-associated Protein 1 Light Chain 3 (LC3) Interacts with Bnip3 Protein to Selectively Remove Endoplasmic Reticulum and Mitochondria via Autophagy *. Journal of Biological Chemistry 2012; 287:19094–104.

16. Pereira RO, Tadinada SM, Zasadny FM, Oliveira KJ, Pires KMP, Olvera A, Jeffers J, Souvenir R, Mcglauflin R, Seei A, et al. OPA1 deficiency promotes secretion of FGF21 from muscle that prevents obesity and insulin resistance. The EMBO Journal 2017; 36:2126–45.

17. Mughal W, Martens M, Field J, Chapman D, Huang J, Rattan S, Hai Y, Cheung KG, Kereliuk S, West AR, et al. Myocardin regulates mitochondrial calcium homeostasis and prevents permeability transition. Cell Death Differ 2018; 25:1732– 48.

18. Trost B, Kindrachuk J, Määttänen P, Napper S, Kusalik A. PIIKA 2: An Expanded, Web-Based Platform for Analysis of Kinome Microarray Data. PLoS ONE 2013; 8:e80837.

19. Kindrachuk J, Falcinelli S, Wada J, Kuhn JH, Hensley LE, Jahrling PB. Systems kinomics for characterizing host responses to high consequence pathogens at the NIH/NIAID Integrated Research Facility-Frederick. Pathogens and Disease 2014; 71:190–8.

20. Bassel-Duby R, Olson EN. Signaling pathways in skeletal muscle remodeling. Annual review of biochemistry 2006; 75:19–37.

21. Grade CVC, Mantovani CS, Alvares LE. Myostatin gene promoter: structure, conservation and importance as a target for muscle modulation. Journal of Animal Science and Biotechnology 2019; 10:32.

22. Poole LP, Bock-Hughes A, Berardi DE, Macleod KF. ULK1 promotes mitophagy via phosphorylation and stabilization of BNIP3. Sci Rep 2021; 11:1–15.

23. Field JT, Martens MD, Mughal W, Hai Y, Chapman D, Hatch GM, Ivanco TL, Diehl-Jones W, Gordon JW. Misoprostol regulates Bnip3 repression and alternative splicing to control cellular calcium homeostasis during hypoxic stress. Cell Death Discovery 2018; 5:37.

24. Mughal W, Nguyen L, Pustylnik S, da Silva Rosa SC, Piotrowski S, Chapman D, Du M, Alli NS, Grigull J, Halayko AJ, et al. A conserved MADS-box phosphorylation motif regulates differentiation and mitochondrial function in skeletal, cardiac, and smooth muscle cells. Cell Death Dis 2015; 6:e1944.

25. Li Y-P. TNF-alpha is a mitogen in skeletal muscle. Am J Physiol Cell Physiol 2003; 285:C370–376.

26. Dick SA, Chang NC, Dumont NA, Bell RAV, Putinski C, Kawabe Y, Litchfield DW, Rudnicki MA, Megeney LA. Caspase 3 cleavage of Pax7 inhibits self-renewal of satellite cells. Proc Natl Acad Sci U S A 2015; 112:E5246–5252.

27. Loboda A, Dulak J. NRF2 and its targets in skeletal muscle repair and regeneration. Antioxid Redox Signal 2023;

28. Palacios D, Mozzetta C, Consalvi S, Caretti G, Saccone V, Proserpio V, Marquez VE, Valente S, Mai A, Forcales SV, et al. TNF/p38α/polycomb signaling to Pax7 locus in satellite cells links inflammation to the epigenetic control of muscle regeneration. Cell Stem Cell 2010; 7:455–69.

29. Peregrin S, Jurado-Pueyo M, Campos PM, Sanz-Moreno V, Ruiz-Gomez A, Crespo P, Mayor F, Murga C. Phosphorylation of p38 by GRK2 at the Docking Groove Unveils a Novel Mechanism for Inactivating p38MAPK. Current Biology 2006; 16:2042–7.

30. Diwan A, Wansapura J, Syed FM, Matkovich SJ, Lorenz JN, Dorn GW. Nix-Mediated Apoptosis Links Myocardial Fibrosis, Cardiac Remodeling, and Hypertrophy Decompensation. Circulation 2008; 117:396–404.

31. Novak I, Kirkin V, Mcewan DG, Zhang J, Wild P, Rozenknop A, Rogov V, Löhr F, Popovic D, Occhipinti A, et al. Nix is a selective autophagy receptor for mitochondrial clearance. EMBO reports 2010; 11:45–51.

32. Samuel VT, Shulman GI. The pathogenesis of insulin resistance: integrating signaling pathways and substrate flux. J Clin Invest 2016; 126:12–22.

33. Wang M, Yu H, Kim YS, Bidwell CA, Kuang S. Myostatin facilitates slow and inhibits fast myosin heavy chain expression during myogenic differentiation. Biochem Biophys Res Commun 2012; 426:83–8.

34. Rodgers BD, Ward CW. Myostatin/Activin Receptor Ligands In Muscle And The Development Status Of Attenuating Drugs. Endocr Rev 2021; :bnab030.

35. Lee S-J. Targeting the myostatin signaling pathway to treat muscle loss and metabolic dysfunction. J Clin Invest [Internet] 2021 [cited 2023 Feb 1]; 131. Available from: https://www.jci.org/articles/view/148372

36. Shintaku J, Guttridge DC. Analysis of Aerobic Respiration in Intact Skeletal Muscle Tissue by Microplate-Based Respirometry. Methods Mol Biol 2016; 1460:337–43.

37. Martens MD, Seshadri N, Nguyen L, Chapman D, Henson ES, Xiang B, Falk L, Mendoza A, Rattan S, Field JT, et al. Misoprostol treatment prevents hypoxia-induced cardiac dysfunction through a 14-3-3 and PKA regulatory motif on Bnip3. Cell Death Dis 2021; 12:1105.

