## Supplemental Material for "The mitophagy receptor BNIP3L/Nix coordinates nuclear calcium signaling to modulate the muscle phenotype"

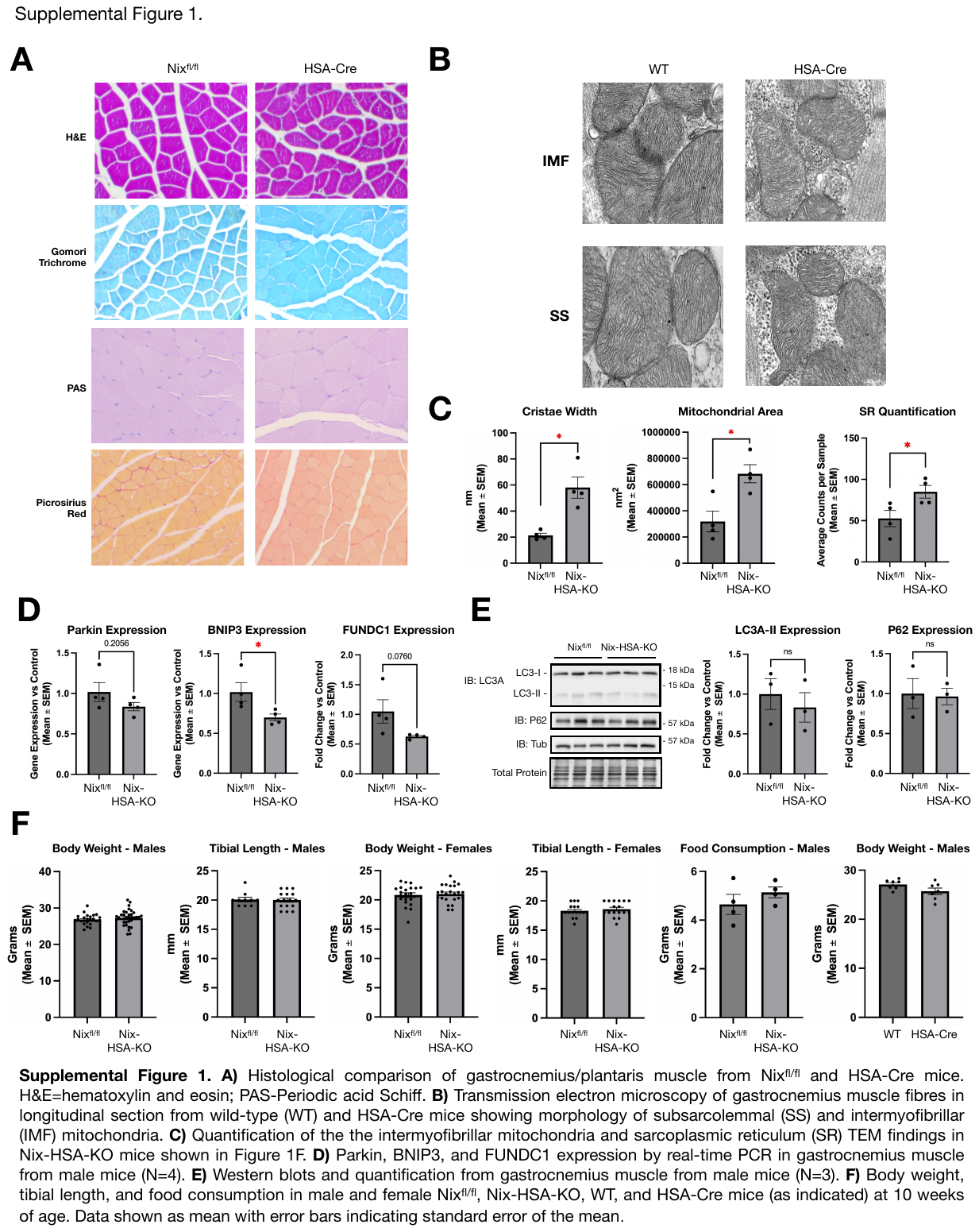

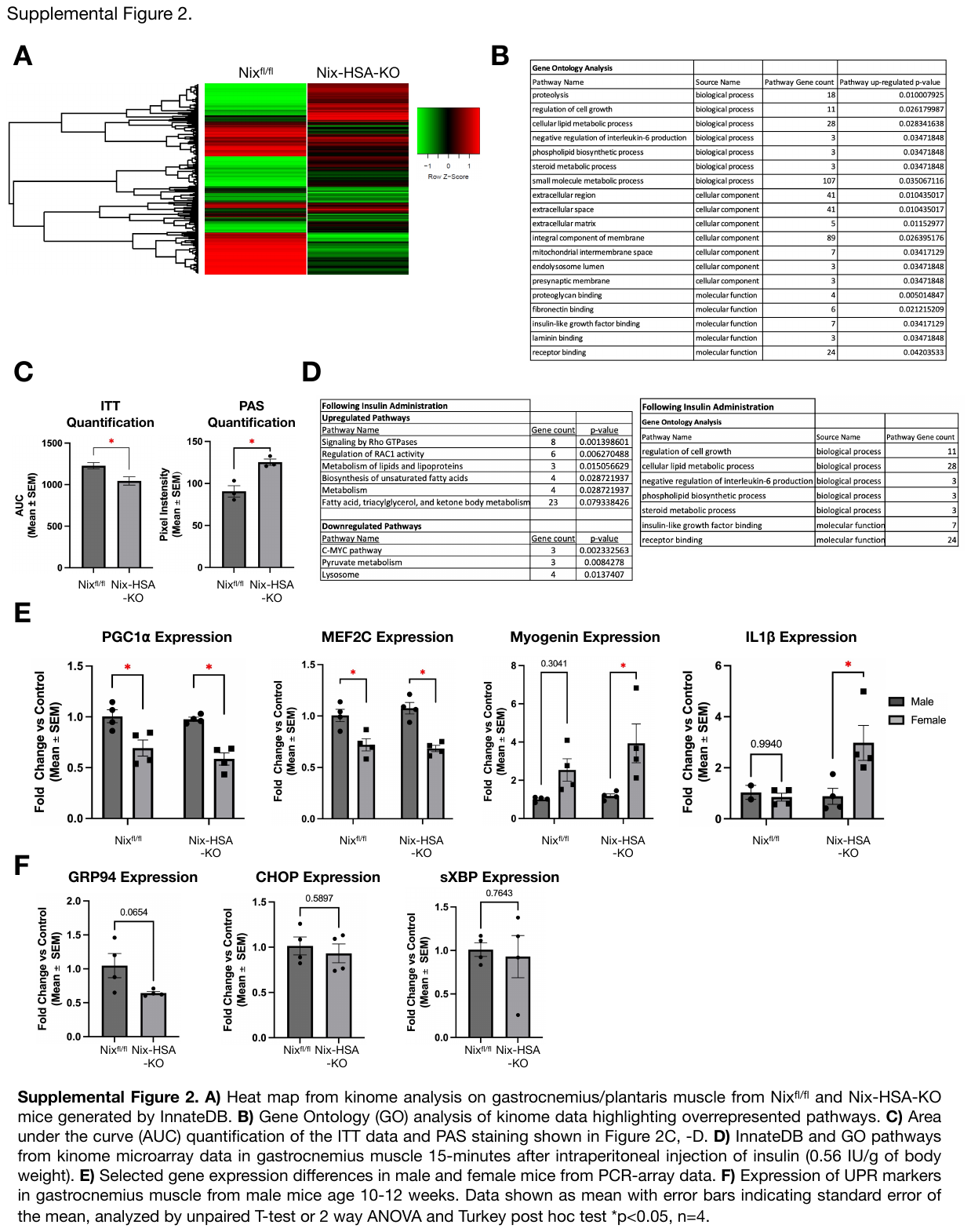

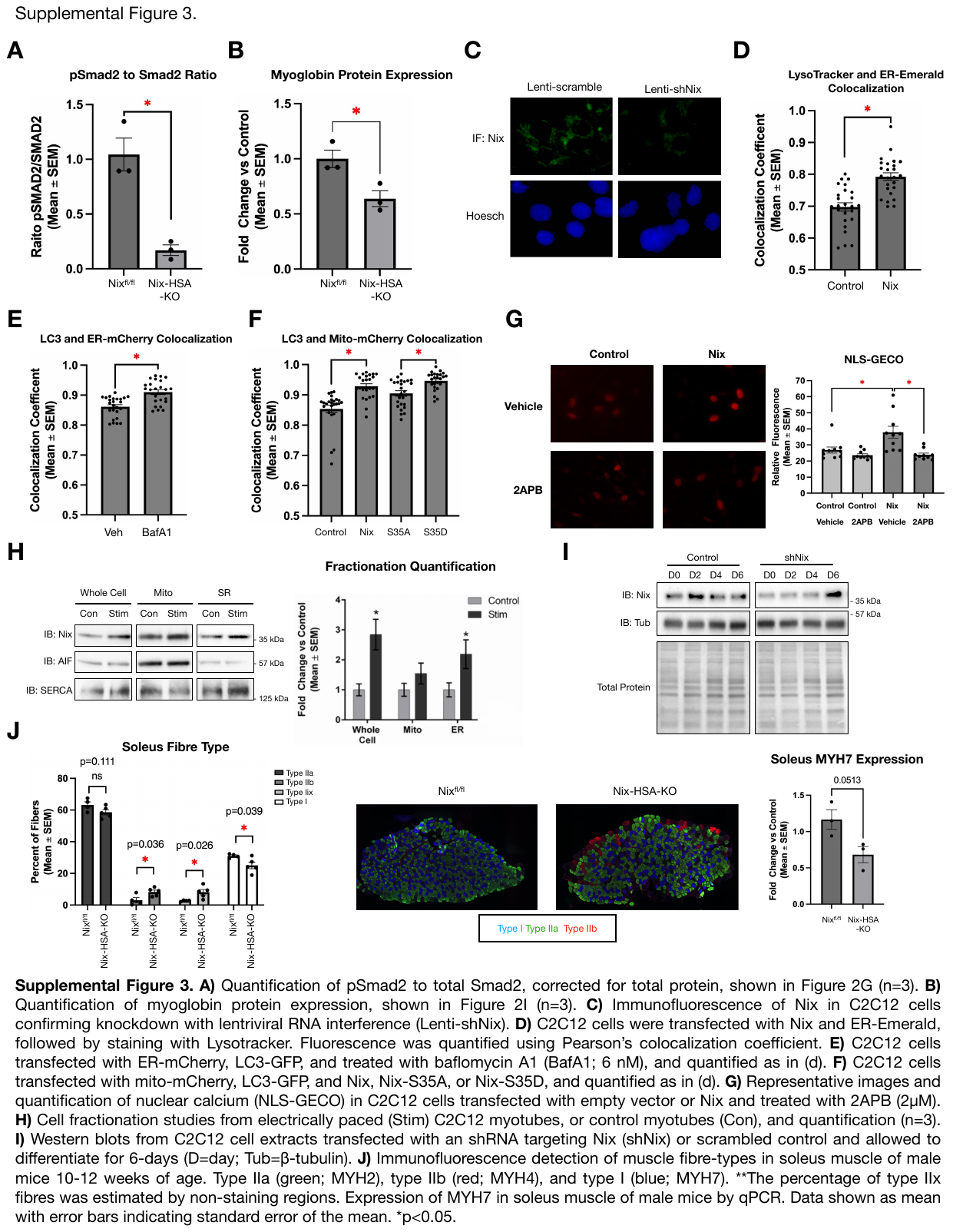

| Supplemental table 1. qPCR array (Myogenesis and Myopathy array) in gastrocnemius/plantaris, soleus, EDL, or male/female as indicated. 2-ΔΔCT indicates fold change vs control (Nix ^fl/fl^). | | | | | | | | | | |
| --- | --- | --- | --- | --- | --- | --- | --- | --- | --- | --- |
|  | **Gastrocnemius/Plantaris** | | | | | | | | | |
|  | **Female** | | | | | **Male** | | | | |
|  | **Nix ^fl/fl^** | | **Nix-HSA-KO** | | | **Nix ^fl/fl^** | | **Nix-HSA-KO** | | |
| **Gene** | **Mean CT** | **ΔCT** | **Mean CT** | **ΔCT** | **2-ΔΔCT** | **Mean CT** | **ΔCT** | **Mean CT** | **ΔCT** | **2-ΔΔCT** |
| Acta1 | 17.157 | -8.292 | 16.235 | -8.295 | 1.002 | 16.691 | -8.337 | 16.700 | -8.410 | 1.052 |
| Akt2 | 25.712 | 0.263 | 24.919 | 0.389 | 0.916 | 24.842 | -0.187 | 25.035 | -0.075 | 0.926 |
| Casp3 | 31.801 | 6.352 | 30.223 | 5.693 | 1.578 | 31.653 | 6.625 | 31.103 | 5.994 | 1.549 |
| Des | 21.165 | -4.284 | 20.173 | -4.357 | 1.051 | 20.273 | -4.755 | 20.459 | -4.651 | 0.930 |
| Gusb | 30.680 | 5.231 | 29.141 | 4.611 | 1.536 | 30.246 | 5.218 | 30.096 | 4.986 | 1.174 |
| Ikbkb | 29.454 | 4.005 | 28.301 | 3.770 | 1.177 | 28.847 | 3.819 | 28.871 | 3.761 | 1.041 |
| Mapk8 | 29.070 | 3.621 | 28.175 | 3.645 | 0.983 | 28.273 | 3.245 | 28.497 | 3.387 | 0.906 |
| Myh1 | 21.761 | -3.688 | 19.888 | -4.642 | 1.938 | 21.133 | -3.895 | 21.392 | -3.718 | 0.884 |
| Pax3 | 35.333 | 9.883 | 35.115 | 10.585 | 0.615 | 34.822 | 9.794 | 37.444 | 12.334 | 0.172 |
| Prkab2 | 25.177 | -0.272 | 24.333 | -0.197 | 0.949 | 24.049 | -0.979 | 24.409 | -0.700 | 0.824 |
| Tnf | 36.868 | 11.419 | 33.013 | 8.483 | 7.652 | 36.250 | 11.222 | 34.197 | 9.087 | 4.392 |
| Atp2a1 | 18.713 | -6.736 | 17.934 | -6.596 | 0.908 | 17.920 | -7.108 | 18.150 | -6.960 | 0.902 |
| Cast | 26.612 | 1.163 | 25.603 | 1.073 | 1.064 | 26.055 | 1.027 | 26.327 | 1.218 | 0.876 |
| Dmd | 27.319 | 1.869 | 27.286 | 2.756 | 0.541 | 26.562 | 1.534 | 27.192 | 2.083 | 0.684 |
| Hdac5 | 27.316 | 1.867 | 26.520 | 1.990 | 0.919 | 27.081 | 2.053 | 27.117 | 2.007 | 1.032 |
| Il1b | 36.644 | 11.195 | 33.938 | 9.408 | 3.451 | 35.914 | 10.886 | 36.425 | 11.315 | 0.743 |
| Mb | 21.197 | -4.252 | 19.656 | -4.875 | 1.539 | 20.586 | -4.442 | 20.830 | -4.280 | 0.893 |
| Myh2 | 22.887 | -2.562 | 21.707 | -2.823 | 1.198 | 22.676 | -2.352 | 22.960 | -2.150 | 0.869 |
| Pax7 | 32.676 | 7.227 | 31.254 | 6.723 | 1.418 | 32.144 | 7.116 | 31.362 | 6.252 | 1.819 |
| Prkag1 | 27.660 | 2.211 | 26.458 | 1.928 | 1.217 | 27.176 | 2.148 | 27.428 | 2.318 | 0.889 |
| Tnnc1 | 25.655 | 0.206 | 24.477 | -0.053 | 1.197 | 25.930 | 0.902 | 25.711 | 0.601 | 1.232 |
| Actn3 | 20.089 | -5.360 | 19.543 | -4.987 | 0.772 | 19.360 | -5.668 | 19.685 | -5.425 | 0.845 |
| B2m | 24.754 | -0.695 | 23.340 | -1.190 | 1.410 | 24.512 | -0.516 | 24.454 | -0.656 | 1.101 |
| Cav1 | 25.721 | 0.272 | 24.650 | 0.120 | 1.111 | 25.290 | 0.262 | 25.198 | 0.088 | 1.128 |
| Dmpk | 25.715 | 0.266 | 23.694 | -0.836 | 2.147 | 24.512 | -0.516 | 24.497 | -0.613 | 1.069 |
| Hk2 | 26.361 | 0.912 | 25.383 | 0.853 | 1.042 | 25.153 | 0.125 | 25.215 | 0.106 | 1.013 |
| Il6 | 36.064 | 10.615 | 34.775 | 10.245 | 1.292 | 35.133 | 10.105 | 35.161 | 10.052 | 1.038 |
| Mef2c | 23.818 | -1.631 | 22.964 | -1.566 | 0.956 | 22.908 | -2.120 | 22.890 | -2.219 | 1.071 |
| Myod1 | 28.530 | 3.081 | 27.273 | 2.743 | 1.264 | 28.301 | 3.273 | 28.222 | 3.112 | 1.118 |
| Pdk4 | 25.309 | -0.140 | 24.594 | 0.064 | 0.868 | 24.302 | -0.726 | 24.253 | -0.856 | 1.094 |
| Prkag3 | 26.178 | 0.729 | 25.519 | 0.988 | 0.836 | 24.873 | -0.155 | 24.948 | -0.162 | 1.005 |
| Tnni2 | 18.439 | -7.010 | 17.987 | -6.544 | 0.724 | 17.905 | -7.123 | 18.010 | -7.100 | 0.984 |
| Acvr2b | 28.687 | 3.238 | 28.010 | 3.480 | 0.846 | 28.180 | 3.152 | 28.198 | 3.088 | 1.046 |
| Bcl2 | 31.478 | 6.028 | 30.445 | 5.915 | 1.082 | 30.977 | 5.949 | 30.867 | 5.758 | 1.142 |
| Cav3 | 26.925 | 1.475 | 25.856 | 1.326 | 1.109 | 26.084 | 1.055 | 26.075 | 0.966 | 1.064 |
| Dysf | 27.912 | 2.463 | 26.699 | 2.169 | 1.226 | 27.039 | 2.011 | 27.026 | 1.917 | 1.068 |
| Hsp90ab1 | 23.378 | -2.071 | 22.596 | -1.934 | 0.909 | 22.823 | -2.205 | 22.904 | -2.206 | 1.001 |
| Lep | 31.999 | 6.550 | 32.494 | 7.964 | 0.375 | 32.881 | 7.853 | 32.303 | 7.194 | 1.579 |
| Mmp9 | 30.916 | 5.467 | 29.916 | 5.386 | 1.058 | 30.764 | 5.736 | 31.050 | 5.941 | 0.868 |
| Myog | 29.318 | 3.869 | 27.790 | 3.259 | 1.526 | 30.130 | 5.102 | 29.989 | 4.880 | 1.166 |
| Pparg | 30.912 | 5.462 | 30.093 | 5.563 | 0.933 | 31.021 | 5.993 | 30.527 | 5.417 | 1.490 |
| Rhoa | 26.255 | 0.806 | 25.346 | 0.816 | 0.993 | 25.686 | 0.658 | 25.624 | 0.514 | 1.105 |
| Tnnt1 | 25.352 | -0.098 | 23.954 | -0.576 | 1.393 | 25.792 | 0.764 | 25.758 | 0.649 | 1.083 |
| Adipoq | 28.128 | 2.678 | 28.164 | 3.634 | 0.516 | 28.959 | 3.931 | 28.312 | 3.202 | 1.657 |
| Bmp4 | 29.902 | 4.453 | 29.120 | 4.590 | 0.909 | 30.144 | 5.116 | 29.928 | 4.819 | 1.229 |
| Cryab | 22.541 | -2.908 | 21.049 | -3.481 | 1.488 | 22.434 | -2.594 | 22.526 | -2.584 | 0.993 |
| Fbxo32 | 25.334 | -0.115 | 24.563 | 0.033 | 0.903 | 24.425 | -0.603 | 24.703 | -0.407 | 0.872 |
| Igf1 | 27.636 | 2.187 | 26.720 | 2.190 | 0.998 | 26.701 | 1.673 | 26.923 | 1.813 | 0.908 |
| Lmna | 26.740 | 1.291 | 25.818 | 1.288 | 1.002 | 26.376 | 1.348 | 26.326 | 1.216 | 1.096 |
| Mstn | 26.796 | 1.347 | 26.502 | 1.972 | 0.648 | 25.631 | 0.603 | 26.254 | 1.144 | 0.687 |
| Myot | 22.731 | -2.718 | 21.846 | -2.684 | 0.977 | 21.959 | -3.069 | 22.169 | -2.941 | 0.915 |
| Ppargc1a | 27.566 | 2.117 | 26.886 | 2.356 | 0.847 | 26.588 | 1.560 | 26.706 | 1.596 | 0.975 |
| Rps6kb1 | 27.882 | 2.432 | 26.741 | 2.211 | 1.166 | 26.975 | 1.947 | 27.213 | 2.103 | 0.897 |
| Tnnt3 | 18.108 | -7.341 | 17.190 | -7.341 | 0.999 | 17.411 | -7.617 | 17.798 | -7.312 | 0.809 |
| Adrb2 | 28.428 | 2.979 | 28.157 | 3.627 | 0.638 | 27.442 | 2.414 | 27.592 | 2.482 | 0.954 |
| Camk2g | 26.353 | 0.903 | 25.595 | 1.065 | 0.894 | 25.753 | 0.725 | 25.879 | 0.769 | 0.970 |
| Cs | 23.824 | -1.625 | 22.608 | -1.923 | 1.229 | 23.148 | -1.880 | 23.165 | -1.945 | 1.046 |
| Fgf2 | 29.927 | 4.478 | 29.240 | 4.710 | 0.851 | 29.880 | 4.852 | 29.687 | 4.577 | 1.210 |
| Igf2 | 28.013 | 2.563 | 26.073 | 1.542 | 2.029 | 28.171 | 3.143 | 27.787 | 2.677 | 1.381 |
| Mapk1 | 26.845 | 1.396 | 25.614 | 1.084 | 1.242 | 25.806 | 0.778 | 25.977 | 0.868 | 0.940 |
| Musk | 28.548 | 3.099 | 27.743 | 3.213 | 0.924 | 26.377 | 1.349 | 26.930 | 1.820 | 0.721 |
| Neb | 21.129 | -4.320 | 20.280 | -4.250 | 0.952 | 20.134 | -4.894 | 20.254 | -4.856 | 0.974 |
| Ppargc1b | 28.635 | 3.186 | 27.608 | 3.078 | 1.078 | 28.261 | 3.233 | 28.316 | 3.207 | 1.018 |
| Sgca | 25.083 | -0.366 | 24.312 | -0.218 | 0.903 | 24.391 | -0.637 | 24.496 | -0.614 | 0.984 |
| Trim63 | 25.816 | 0.367 | 25.627 | 1.096 | 0.603 | 25.432 | 0.404 | 25.479 | 0.370 | 1.024 |
| Agrn | 30.366 | 4.917 | 30.072 | 5.542 | 0.649 | 30.340 | 5.312 | 30.135 | 5.025 | 1.220 |
| Capn2 | 26.688 | 1.238 | 26.186 | 1.655 | 0.749 | 26.541 | 1.513 | 26.637 | 1.527 | 0.990 |
| Ctnnb1 | 24.892 | -0.557 | 24.401 | -0.129 | 0.743 | 25.014 | -0.014 | 25.115 | 0.005 | 0.987 |
| Foxo1 | 29.218 | 3.768 | 29.259 | 4.729 | 0.514 | 29.281 | 4.253 | 29.261 | 4.152 | 1.072 |
| Igfbp3 | 29.707 | 4.258 | 29.343 | 4.813 | 0.681 | 29.896 | 4.868 | 30.091 | 4.982 | 0.924 |
| Mapk14 | 25.910 | 0.461 | 25.474 | 0.944 | 0.716 | 25.487 | 0.459 | 25.758 | 0.648 | 0.877 |
| Myf5 | 31.743 | 6.294 | 31.668 | 7.138 | 0.557 | 32.139 | 7.111 | 31.990 | 6.880 | 1.174 |
| Nfkb1 | 28.515 | 3.066 | 28.034 | 3.504 | 0.738 | 28.263 | 3.235 | 28.278 | 3.168 | 1.048 |
| Ppp3ca | 24.935 | -0.514 | 24.945 | 0.415 | 0.525 | 25.047 | 0.019 | 25.296 | 0.186 | 0.891 |
| Slc2a4 | 24.013 | -1.436 | 23.791 | -0.739 | 0.617 | 23.927 | -1.101 | 23.824 | -1.285 | 1.137 |
| Ttn | 20.148 | -5.301 | 19.880 | -4.651 | 0.637 | 19.805 | -5.223 | 19.793 | -5.317 | 1.067 |
| Akt1 | 27.593 | 2.144 | 26.969 | 2.439 | 0.815 | 27.432 | 2.404 | 27.367 | 2.257 | 1.107 |
| Capn3 | 26.173 | 0.724 | 25.528 | 0.998 | 0.827 | 25.418 | 0.390 | 25.709 | 0.599 | 0.865 |
| Dag1 | 25.454 | 0.005 | 24.589 | 0.059 | 0.963 | 24.830 | -0.198 | 25.025 | -0.085 | 0.925 |
| Foxo3 | 28.615 | 3.166 | 28.158 | 3.627 | 0.726 | 28.352 | 3.324 | 28.218 | 3.108 | 1.161 |
| Igfbp5 | 24.661 | -0.788 | 23.172 | -1.358 | 1.485 | 23.250 | -1.779 | 23.729 | -1.380 | 0.759 |
| Mapk3 | 28.241 | 2.792 | 27.435 | 2.905 | 0.925 | 28.254 | 3.226 | 28.087 | 2.978 | 1.188 |
| Myf6 | 26.304 | 0.855 | 25.485 | 0.955 | 0.933 | 25.623 | 0.595 | 25.682 | 0.572 | 1.016 |
| Nos2 | 32.101 | 6.652 | 30.034 | 5.504 | 2.216 | 31.037 | 6.009 | 31.083 | 5.973 | 1.025 |
| Prkaa1 | 29.034 | 3.585 | 28.128 | 3.598 | 0.991 | 28.511 | 3.483 | 28.567 | 3.457 | 1.018 |
| Tgfb1 | 30.057 | 4.608 | 29.003 | 4.473 | 1.098 | 29.851 | 4.823 | 29.507 | 4.397 | 1.343 |
| Utrn | 28.340 | 2.891 | 27.102 | 2.571 | 1.248 | 27.651 | 2.623 | 27.598 | 2.488 | 1.098 |
| Other Genes Analyzed | | |  |  |  |  |  |  |  |  |
| GDF15 | ND |  | ND |  |  | 38.37175 | 14.84675 | 38.08575 | 14.7975 | 1.083641 |
| FGF21 | 36.78875 | 8.364 | 35.499 | 7.875 | 1.442286 | 35.89275 | 12.36775 | 35.892 | 12.60375 | 0.884178 |
| *ND= not detected | |  |  |  |  |  |  |  |  |  |

|  | **Soleus** | | | | | **EDL** | | | | |
| --- | --- | --- | --- | --- | --- | --- | --- | --- | --- | --- |
|  | **Nix ^fl/fl^** | | **Nix-HSA-KO** | | | **Nix ^fl/fl^** | | **Nix-HSA-KO** | | |
| **Gene** | **Mean** | **ΔCT** | **Mean** | **ΔCT** | **2-ΔΔCT** | **Mean** | **ΔCT** | **Mean** | **ΔCT** | **2-ΔΔCT** |
| Acta1 | 16.652 | -8.414 | 17.132 | -8.086 | 0.797 | 17.000 | -7.966 | 17.117 | -7.860 | 0.929 |
| Akt2 | 25.051 | -0.015 | 25.478 | 0.259 | 0.827 | 24.863 | -0.104 | 24.967 | -0.010 | 0.937 |
| Casp3 | 30.716 | 5.650 | 30.914 | 5.696 | 0.969 | 31.612 | 6.646 | 31.813 | 6.836 | 0.876 |
| Des | 20.080 | -4.986 | 20.439 | -4.780 | 0.867 | 20.473 | -4.494 | 20.572 | -4.405 | 0.940 |
| Gusb | 29.499 | 4.433 | 29.756 | 4.538 | 0.930 | 30.153 | 5.186 | 30.115 | 5.138 | 1.034 |
| Ikbkb | 28.265 | 3.200 | 28.529 | 3.311 | 0.926 | 28.875 | 3.909 | 28.751 | 3.774 | 1.098 |
| Mapk8 | 27.469 | 2.403 | 28.107 | 2.889 | 0.714 | 28.097 | 3.130 | 28.298 | 3.321 | 0.876 |
| Myh1 | 19.845 | -5.221 | 20.638 | -4.581 | 0.641 | 19.447 | -5.519 | 19.595 | -5.382 | 0.909 |
| Pax3 | 39.339 | 14.273 | ND | ND |  | 35.284 | 10.318 | 35.445 | 10.468 | 0.901 |
| Prkab2 | 26.849 | 1.783 | 27.570 | 2.352 | 0.674 | 24.477 | -0.489 | 24.916 | -0.061 | 0.743 |
| Tnf | 34.037 | 8.971 | 34.368 | 9.149 | 0.884 | 35.749 | 10.783 | 35.135 | 10.158 | 1.542 |
| Atp2a1 | 19.287 | -5.779 | 19.706 | -5.513 | 0.831 | 18.112 | -6.854 | 18.309 | -6.668 | 0.879 |
| Cast | 25.101 | 0.035 | 25.858 | 0.639 | 0.658 | 25.706 | 0.740 | 25.903 | 0.926 | 0.879 |
| Dmd | 25.828 | 0.762 | 26.649 | 1.430 | 0.629 | 26.274 | 1.308 | 26.678 | 1.701 | 0.762 |
| Hdac5 | 26.357 | 1.291 | 26.731 | 1.512 | 0.858 | 26.788 | 1.822 | 26.863 | 1.886 | 0.956 |
| Il1b | 35.300 | 10.234 | 34.637 | 9.418 | 1.761 | 35.777 | 10.811 | 36.087 | 11.110 | 0.813 |
| Mb | 17.551 | -7.514 | 18.355 | -6.863 | 0.637 | 19.483 | -5.484 | 19.625 | -5.352 | 0.913 |
| Myh2 | 19.334 | -5.732 | 19.943 | -5.275 | 0.729 | 21.798 | -3.168 | 22.162 | -2.815 | 0.783 |
| Pax7 | 30.934 | 5.869 | 30.839 | 5.621 | 1.187 | 31.907 | 6.941 | 31.459 | 6.481 | 1.375 |
| Prkag1 | 26.246 | 1.180 | 26.949 | 1.731 | 0.683 | 27.009 | 2.043 | 27.153 | 2.176 | 0.912 |
| Tnnc1 | 19.977 | -5.089 | 20.479 | -4.740 | 0.785 | 27.197 | 2.230 | 27.056 | 2.079 | 1.111 |
| Actn3 | 25.692 | 0.626 | 24.854 | -0.365 | 1.987 | 20.200 | -4.767 | 20.396 | -4.581 | 0.879 |
| B2m | 23.708 | -1.358 | 23.843 | -1.375 | 1.012 | 24.533 | -0.433 | 24.377 | -0.600 | 1.123 |
| Cav1 | 24.427 | -0.639 | 24.634 | -0.585 | 0.963 | 25.297 | 0.330 | 25.211 | 0.234 | 1.069 |
| Dmpk | 23.800 | -1.265 | 24.414 | -0.804 | 0.726 | 24.540 | -0.427 | 24.591 | -0.386 | 0.972 |
| Hk2 | 24.750 | -0.316 | 25.026 | -0.192 | 0.918 | 24.992 | 0.026 | 25.127 | 0.150 | 0.918 |
| Il6 | 35.032 | 9.967 | 35.691 | 10.473 | 0.704 | 34.953 | 9.986 | 35.736 | 10.759 | 0.585 |
| Mef2c | 22.359 | -2.706 | 22.859 | -2.359 | 0.786 | 22.863 | -2.103 | 23.045 | -1.932 | 0.889 |
| Myod1 | 29.039 | 3.974 | 28.924 | 3.706 | 1.204 | 27.923 | 2.957 | 28.162 | 3.185 | 0.854 |
| Pdk4 | 22.199 | -2.867 | 22.523 | -2.696 | 0.888 | 24.389 | -0.577 | 24.350 | -0.627 | 1.035 |
| Prkag3 | 29.292 | 4.227 | 29.131 | 3.912 | 1.244 | 25.885 | 0.918 | 26.276 | 1.299 | 0.768 |
| Tnni2 | 18.726 | -6.339 | 18.829 | -6.390 | 1.035 | 17.880 | -7.087 | 17.981 | -6.996 | 0.939 |
| Acvr2b | 28.624 | 3.558 | 28.888 | 3.670 | 0.926 | 28.087 | 3.121 | 28.258 | 3.281 | 0.895 |
| Bcl2 | 30.972 | 5.906 | 31.084 | 5.866 | 1.029 | 31.131 | 6.165 | 31.035 | 6.058 | 1.077 |
| Cav3 | 25.983 | 0.917 | 26.072 | 0.853 | 1.045 | 26.106 | 1.140 | 26.065 | 1.088 | 1.036 |
| Dysf | 27.205 | 2.139 | 27.417 | 2.199 | 0.959 | 27.063 | 2.096 | 27.214 | 2.237 | 0.907 |
| Hsp90ab1 | 22.446 | -2.620 | 22.847 | -2.372 | 0.842 | 22.774 | -2.192 | 23.023 | -1.954 | 0.848 |
| Lep | 32.098 | 7.032 | 33.372 | 8.153 | 0.460 | 33.978 | 9.011 | 34.537 | 9.559 | 0.684 |
| Mmp9 | 33.500 | 8.434 | 33.294 | 8.076 | 1.282 | 31.226 | 6.260 | 31.397 | 6.420 | 0.895 |
| Myog | 29.330 | 4.264 | 29.944 | 4.726 | 0.726 | 30.701 | 5.735 | 30.771 | 5.794 | 0.960 |
| Pparg | 29.540 | 4.474 | 29.520 | 4.302 | 1.127 | 30.792 | 5.826 | 30.908 | 5.931 | 0.930 |
| Rhoa | 25.338 | 0.272 | 25.384 | 0.165 | 1.077 | 25.812 | 0.846 | 25.844 | 0.867 | 0.986 |
| Tnnt1 | 19.848 | -5.218 | 20.603 | -4.616 | 0.659 | 26.939 | 1.972 | 26.935 | 1.958 | 1.010 |
| Adipoq | 28.244 | 3.178 | 28.385 | 3.167 | 1.008 | 29.433 | 4.467 | 29.780 | 4.802 | 0.792 |
| Bmp4 | 29.884 | 4.818 | 29.821 | 4.603 | 1.161 | 30.116 | 5.150 | 30.115 | 5.138 | 1.008 |
| Cryab | 19.704 | -5.362 | 20.260 | -4.959 | 0.756 | 21.923 | -3.043 | 22.126 | -2.851 | 0.875 |
| Fbxo32 | 24.387 | -0.679 | 24.804 | -0.414 | 0.832 | 24.629 | -0.338 | 24.788 | -0.189 | 0.902 |
| Igf1 | 27.331 | 2.265 | 27.804 | 2.585 | 0.801 | 27.073 | 2.107 | 27.040 | 2.063 | 1.031 |
| Lmna | 25.313 | 0.247 | 25.602 | 0.384 | 0.910 | 26.159 | 1.192 | 26.147 | 1.170 | 1.016 |
| Mstn | 31.567 | 6.501 | 31.676 | 6.458 | 1.030 | 26.034 | 1.068 | 26.477 | 1.500 | 0.741 |
| Myot | 20.857 | -4.209 | 21.327 | -3.892 | 0.802 | 21.594 | -3.372 | 21.819 | -3.158 | 0.862 |
| Ppargc1a | 25.658 | 0.592 | 26.037 | 0.818 | 0.855 | 26.319 | 1.353 | 26.407 | 1.430 | 0.948 |
| Rps6kb1 | 26.809 | 1.744 | 27.462 | 2.243 | 0.707 | 27.060 | 2.093 | 27.161 | 2.184 | 0.939 |
| Tnnt3 | 18.332 | -6.734 | 19.023 | -6.195 | 0.689 | 17.678 | -7.288 | 17.885 | -7.092 | 0.873 |
| Adrb2 | 27.782 | 2.716 | 27.782 | 2.563 | 1.112 | 27.630 | 2.663 | 27.892 | 2.915 | 0.840 |
| Camk2g | 26.710 | 1.645 | 26.873 | 1.654 | 0.993 | 25.856 | 0.889 | 26.084 | 1.107 | 0.860 |
| Cs | 22.528 | -2.538 | 22.921 | -2.298 | 0.847 | 23.158 | -1.808 | 23.285 | -1.692 | 0.922 |
| Fgf2 | 29.012 | 3.946 | 29.137 | 3.919 | 1.019 | 29.635 | 4.668 | 29.779 | 4.802 | 0.912 |
| Igf2 | 27.902 | 2.836 | 27.800 | 2.581 | 1.193 | 28.461 | 3.495 | 28.478 | 3.501 | 0.995 |
| Mapk1 | 25.456 | 0.390 | 25.927 | 0.709 | 0.802 | 25.794 | 0.828 | 26.003 | 1.026 | 0.871 |
| Musk | 26.121 | 1.055 | 26.540 | 1.322 | 0.831 | 26.758 | 1.791 | 27.290 | 2.313 | 0.696 |
| Neb | 19.942 | -5.124 | 20.477 | -4.742 | 0.767 | 19.990 | -4.976 | 20.293 | -4.684 | 0.817 |
| Ppargc1b | 27.062 | 1.997 | 27.379 | 2.161 | 0.892 | 28.067 | 3.101 | 28.086 | 3.109 | 0.995 |
| Sgca | 24.427 | -0.639 | 24.697 | -0.521 | 0.921 | 24.327 | -0.640 | 24.544 | -0.433 | 0.866 |
| Trim63 | 25.882 | 0.817 | 26.040 | 0.821 | 0.997 | 25.982 | 1.016 | 26.154 | 1.177 | 0.894 |
| Agrn | 29.743 | 4.677 | 29.322 | 4.104 | 1.488 | 30.097 | 5.131 | 30.021 | 5.044 | 1.062 |
| Capn2 | 26.378 | 1.313 | 26.634 | 1.416 | 0.931 | 26.632 | 1.666 | 26.688 | 1.711 | 0.969 |
| Ctnnb1 | 24.475 | -0.591 | 25.025 | -0.194 | 0.759 | 24.982 | 0.015 | 25.046 | 0.068 | 0.964 |
| Foxo1 | 28.885 | 3.819 | 29.090 | 3.872 | 0.964 | 29.347 | 4.380 | 29.479 | 4.502 | 0.919 |
| Igfbp3 | 29.678 | 4.612 | 29.393 | 4.175 | 1.354 | 29.995 | 5.028 | 30.087 | 5.110 | 0.945 |
| Mapk14 | 26.283 | 1.217 | 26.578 | 1.359 | 0.906 | 25.997 | 1.031 | 26.150 | 1.172 | 0.906 |
| Myf5 | 30.934 | 5.868 | 31.025 | 5.807 | 1.043 | 31.800 | 6.834 | 32.044 | 7.067 | 0.851 |
| Nfkb1 | 28.067 | 3.002 | 28.272 | 3.053 | 0.965 | 28.258 | 3.292 | 28.491 | 3.514 | 0.857 |
| Ppp3ca | 25.762 | 0.696 | 26.197 | 0.978 | 0.823 | 25.052 | 0.086 | 25.303 | 0.326 | 0.847 |
| Slc2a4 | 24.001 | -1.065 | 24.021 | -1.198 | 1.097 | 23.950 | -1.016 | 24.053 | -0.924 | 0.938 |
| Ttn | 19.257 | -5.809 | 19.669 | -5.550 | 0.836 | 19.445 | -5.521 | 19.714 | -5.263 | 0.836 |
| Akt1 | 26.521 | 1.455 | 26.661 | 1.443 | 1.008 | 27.130 | 2.164 | 27.269 | 2.292 | 0.915 |
| Capn3 | 25.344 | 0.278 | 25.677 | 0.458 | 0.883 | 25.475 | 0.509 | 25.740 | 0.763 | 0.839 |
| Dag1 | 24.696 | -0.369 | 24.898 | -0.320 | 0.966 | 25.021 | 0.054 | 25.205 | 0.228 | 0.887 |
| Foxo3 | 28.134 | 3.069 | 28.127 | 2.908 | 1.118 | 28.194 | 3.228 | 28.480 | 3.503 | 0.827 |
| Igfbp5 | 24.631 | -0.435 | 25.309 | 0.091 | 0.694 | 23.237 | -1.729 | 23.764 | -1.213 | 0.699 |
| Mapk3 | 27.583 | 2.517 | 27.550 | 2.331 | 1.138 | 28.106 | 3.139 | 28.086 | 3.109 | 1.021 |
| Myf6 | 25.870 | 0.804 | 26.168 | 0.950 | 0.904 | 25.626 | 0.660 | 25.624 | 0.647 | 1.009 |
| Nos2 | 30.178 | 5.112 | 30.368 | 5.149 | 0.975 | 30.950 | 5.983 | 30.950 | 5.973 | 1.007 |
| Prkaa1 | 27.753 | 2.687 | 28.081 | 2.863 | 0.886 | 28.354 | 3.388 | 28.365 | 3.388 | 1.000 |
| Tgfb1 | 29.240 | 4.174 | 29.067 | 3.849 | 1.253 | 29.875 | 4.909 | 29.871 | 4.894 | 1.010 |
| Utrn | 26.739 | 1.673 | 26.938 | 1.719 | 0.969 | 27.404 | 2.437 | 27.459 | 2.482 | 0.969 |

**Supplemental Materials**

Table of Antibodies:

|  | **Immunofluorescence** | | | | | | |
| --- | --- | --- | --- | --- | --- | --- | --- |
|  | | **Primary Antibody** | **Dilution** |  | | **Secondary** | **Dilution** |
|  | | MYH type IIa (DSHB, SC-71) | 1:600 |  | | Alexa Fluor 488 IgG_1_ | 1:500 |
|  | | MYH Type I (DSHB, BA-D5) | 1:50 |  | | Alexa Fluor 555 IgM | 1:500 |
|  | | MYH Type IIb (DSHB, BF-F3) | 1:100 |  | | Alexa Fluor 350 IgG_2b_ | 1:500 |
|  | | Pax7 (DSHB, PAX7) | 1:10 |  | | Alexa Fluor 488 IgG | 1:500 |
|  | | Laminin (Abcam, ab11575) | 1:500 |  | | Alexa Fluor 568 IgG | 1:500 |
|  | **Immunoblotting** | | | | | | |
|  | | **Antibody** | **Dilution** |  | **Notes** | | |
|  | | Nix (CS, 12396) | 1:1000 |  |  | | |
|  | | Bnip3 (CS, 3769) | 1:1000 |  |  | | |
|  | | Cre (CS, 15036) | 1:1000 |  |  | | |
|  | | β-Tubulin (CS, 86298) | 1:10000 |  |  | | |
|  | | Smad2/3 (CS# 8685) | 1:1000 |  |  | | |
|  | | Phoshp-Smad2 (CS, 18338) | 1:1000 |  |  | | |
|  | | Myoglobin (CS, 25919) | 1:10000 |  | Reduced Protein (5 ug). Reduced Secondary Antibody (1:100000) | | |
|  | | PGC1-α (SC, 13067) | 1:100 |  |  | | |
|  | | NRF2 (CS, 12721) | 1:1000 |  |  | | |
|  | | AIF (CS, 5318 | 1:1000 |  |  | | |
|  | | SERCA CS, 9580) | 1:1000 |  |  | | |
|  | | SQSTM1/P62 (CS, 5114) | 1:1000 |  |  | | |
|  | | LC3A (CS, 4599) | 1:1000 |  |  | | |
|  | | Donkey Anti-Rabbit HRP  (Jackson Immuno #711-035-152) | 1:10000 |  |  | | |
|  | | Donkey Anti-Mouse HRP  (Jackson Immuno #715-035-150) | 1:10000 |  |  | | |
| All DSBH antibodies used are purchased as concentrated supernatant.  DSHB, Developmental Studies Hybridoma Bank; CS, Cell Signaling; SC, Santa Cruz Biotechnology. | | | | | | | |

Table of Plasmids:

| **Plasmid** | **Addgene Number** | **A Gift From** |
| --- | --- | --- |
| Myc-Nix | #100795 |  |
| Bnip3L RNAi pSuper (shNix) | #17469 | Wafik El-Deiry |
| pLKO-shNix | #100770 |  |
| NFAT-YFP |  | T. Miyake and J. McDermott |
| GW1-Mito-pHRed | #31474 | Gary Yellen |
| CMV-NLS-R-GECO | #32462 | Robert Campbell |
| Myc-Nix-S35A | #197562 |  |
| Myc-Nix-S35D | #197563 |  |
| pEGFP-LC3 | #24920 | Toren Finkel |
| mCherry-ER-3 | #55041 | Michael Davidson |
| mCherry-mito-7 | #55102 | Michael Davidson |
| mEmerald-ER-3 | #54082 | Michael Davidson |
| CMV-ER-LAR-GECO1 | #61244 | Robert Campbell |

Table of Primers: (Not from Myogenesis & Myopathy PCR Array)

| **Gene** | **Primer Sequence** | **Notes** |
| --- | --- | --- |
| Nix | **Forward** TGATGTTGAGATGCACACCAG |  |
|  | **Reverse** GTGGGATGTTTTCGGGTCTA |  |
| Nix 5’ LoxP site | **Forward** GGAGAGACACACATCTGTAGAATAG | Genotyping |
|  | **Reverse** ATGACTTAGCAACATCATACAGTTC |  |
| Nix 3’ LoxP site | **Forward** AGTCAGGGCTGTATAGTAAGGT | Genotyping |
|  | **Reverse** GGTGTATATGTGGAAGCCAGAG |  |
| HSA-Cre | **Forward** GCGGTCTGGCAGTAAAAACTATC | Genotyping |
|  | **Reverse** GTGAAACAGCATTGCTGTCACTT |  |
| MYH1 | **Forward** CACTTACCAAACTGAGGAAGACC |  |
|  | **Reverse** CCAGGTTGACGTTGGATTG |  |
| MYH2 | **Forward** GAGTGAGCTGAAGTCGAAGGA |  |
|  | **Reverse** CCCCTTGATAACTGAGAAACCAG |  |
| MYH3 | **Forward** GCTAACACGGGAGAAGAAGG |  |
|  | **Reverse** TTTTGTTCCTTTCTAGGTCCACA |  |
| MYH4 | **Forward** TAGGAACACACAGGGAATGCT |  |
|  | **Reverse** TAGCTCTTGCTCAGCCACTC |  |
| MYH7 | **Forward** AAGAGCCGGGACATTGGT |  |
|  | **Reverse** TTGGAGCTGGGTAGCACAAG |  |
| FUNDC1 | **Forward** CGGACCTATGGTAGAAAAATACTCA  **Reverse** AGAAGGAAACCACCACCTACTG |  |
| CHOP | **Forward** AGCCTGGTATGAGGATCTGC  **Reverse** ACGCAGGGTCAAGAGTAGTG |  |
| GRP94 | **Forward** AGAATGAAGGAAAAACAGGACAA  **Reverse** TCAGAAGTCTCTCAACAAATGGA |  |
| XBP1 | **Forward** CTGAGTCCGCAGCAGGTG  **Reverse** AGAGTCCATGGGAAGATGTTCTG | For splice variant |
| BNIP3 | **Forward** CCAGACACCACAAGATACCAAC  **Reverse** GTCGACTTGACCAATCCCATATC |  |
| FGF21 | **Forward** CACAGATGACGACCAAGACAC  **Reverse** GACACCCAGGATTTGAATGACC |  |
| GDF15 | **Forward** AGGACTCGAACTCAGAACCAA  **Reverse** CTTCAGGGGCCTAGTGATGT |  |
| PARKIN | **Forward** GCTCAAGGAAGTGGTTGCTA  **Reverse** ATGACTTCTCCTCCGTGGTC |  |
| Myoglobin | **Forward** CCAGCCTCTAGCCCAATCA  **Reverse** CCCGGAATGTCTCTTCTTCAG |  |
| L13 | **Forward** AGGAGGCGAAACAAATCCAC  **Reverse** TATGAGCTTGGAGCGGTACTC |  |
| Beta Actin | **Forward** CTGTGTGGATTGGTGGCTCTA  **Reverse** AAAACGCAGCTCAGTAACAGTCC |  |
| Nix-S35A | **Forward:** CCTCAACAGTgccTGGGTGGAGCTACCCATGAACAG | Mutagenesis |
|  | **Reverse:** CCGGCCGGCGGGGGCAGA |  |
| Nix-S35D | **Forward:** CCTCAACAGTgacTGGGTGGAGCTACCCATGAACAGCAGCAATGGC  **Reverse:** CCGGCCGGCGGGGGCAGA | Mutagenesis |
